# Kidney organoids: A system to study human basement membrane assembly in health and disease

**DOI:** 10.1101/2021.06.27.450067

**Authors:** Mychel Raony Paiva Teixeira Morais, Pinyuan Tian, Craig Lawless, Syed Murtuza-Baker, Louise Hopkinson, Steven Woods, Aleksandr Mironov, David Andrew Long, Daniel Gale, Telma Maria Tenorio Zorn, Susan Kimber, Roy Zent, Rachel Lennon

## Abstract

Basement membranes (BMs) are complex macromolecular networks underlying all continuous layers of cells. Essential components include type IV collagen and laminins, which are affected by human genetic defects leading to a range of debilitating conditions including kidney, muscle, and cerebrovascular phenotypes. We investigated the dynamics of BM assembly in human pluripotent stem cell-derived kidney organoids. We resolved their global BM composition and discovered a conserved temporal sequence in BM assembly that paralleled mammalian fetal kidneys. We identified the emergence of key BM isoforms, which were altered by a pathogenic variant in *COL4A5*. Integrating organoid, fetal and adult kidney proteomes we found dynamic regulation of BM composition through development to adulthood, and with single-cell transcriptomic analysis we mapped the cellular origins of BM components. Overall, we define the complex and dynamic nature of vertebrate BM assembly and provide a platform for understanding its wider relevance in human development and disease.

## INTRODUCTION

Basement membranes (BMs) surround tissues providing cells with an interface for physical and signaling interactions (Jayadev and Sherwood, 2017). They are composed of laminins, type IV collagen, nidogens, heparan-sulfate proteoglycans (Kruegel and Miosge, 2010) and many minor components that combine to form biochemically distinct BMs across different tissues (Randles et al., 2017). BMs play active morphogenic roles that are critical for tissue and cell fate specification (Kyprianou et al., 2020; Li et al., 2003), and variants in BM genes are associated with a broad range of human diseases (Chew and Lennon, 2018; Gatseva et al., 2019). Despite increasing knowledge of BM composition and function there is limited understanding about BM assembly, yet this is required for new mechanistic insights into BM-associated human disease.

BMs form early in embryogenesis through binding interactions with cell surface receptors (Miner and Yurchenco, 2004) and typically an initial laminin network is required for further incorporation of collagen IV, nidogen and perlecan into nascent BMs (Jayadev et al., 2019; Matsubayashi et al., 2017) thus following an assembly hierarchy for *de novo* BM formation. BMs are also highly dynamic, remodeling during morphogenesis to form tissue-specific BMs (Bonnans et al., 2014), such as the glomerular basement membrane (GBM) in the kidney, which functions as a size selective filter. Situated between podocytes and endothelial cells in the glomerular capillary wall, the GBM is formed by the fusion of separate podocyte and endothelial BMs, and further remodeled into a mature GBM. This involves replacement of laminin α1β1γ1 (termed laminin-111) and type IV collagen α1α1α2 networks by laminin-511 then −521, and type collagen IV α3α4α5 (Abrahamson and St John, 1993; Abrahamson et al., 2013). These transitions are important for long term GBM function and genetic defects in *COL4A3, COL4A4* and *COL4A5* or the laminin gene *LAMB2* cause defective GBMs and human disease (Barker et al., 1990; Zenker et al., 2004).

The study of BM assembly is challenging due to the technical difficulties in tracking large, spatiotemporally regulated components. Most understanding about vertebrate BMs comes from immunolocalization and knock-out studies (Abrahamson et al., 2013) and for composition, mass spectrometry (MS)-based proteomics has enabled global analysis (Naba et al., 2016; Randles et al., 2015). Proteomics also allows time course studies, which have provided insight into matrix dynamics during development and in disease progression (Hebert et al., 2020; Lipp et al., 2021; Naba et al., 2017). However, proteomics lacks the spatial context that is captured by localization studies including fluorescent tagging of endogenous proteins. Such investigations in *Drosophila* and *C. elegans* have unraveled dynamic features of BM assembly in embryogenesis and repair (Howard et al., 2019; Keeley et al., 2020; Matsubayashi et al., 2020). The development of a system to study vertebrate BM assembly would facilitate investigation of both morphogenesis and disease.

Kidney organoids generated from pluripotent stem cells (PSCs) contain self-organised 3D structures with multiple kidney cell types and they represent an attractive system for investigating early development (Combes et al., 2019a; Takasato et al., 2015). Organoids derived from induced PSCs (iPSCs), reprogrammed from patient somatic cells have further utility in personalized disease modelling and therapy screening (Czerniecki et al., 2018; Forbes et al., 2018). The nephron is the functional unit of the kidney and during differentiation, kidney organoids pattern into early nephron structures with clusters of podocytes and endothelial cells, and a complex tubular epithelial system. Furthermore organoids show transcriptomic homology to the first trimester human fetal kidney (Takasato et al., 2015) and differentiation is further advanced by *in vivo* implantation (Bantounas et al., 2018). Whilst understanding about cell types in kidney organoids has advanced significantly, there is a knowledge gap about extracellular matrix and BM assembly during differentiation.

We investigated BM assembly during kidney development using organoids and fetal kidney tissue. With proteomics we defined a complex sequence of BM assembly during organoid differentiation and demonstrated the utility of this experimental system for investigating BM remodelling in both early development and human disease. Furthermore, we relate the organoid matrix to the E19 mouse kidney and adult human kidney matrix and define the cellular origins of BM components. Overall, we demonstrate that organoids represent a high-fidelity system to study the dynamics of human BM assembly.

## RESULTS

### Kidney organoids form BM networks that are altered with defective *COL4A5*

To improve understanding of BM assembly and regulation, we investigated human kidney organoids. We differentiated wild type iPSCs into intermediate mesoderm cells in 2D culture, and then 3D-kidney organoids (**Figures 1A** and **S1A**). We confirmed differentiation to glomerular clusters (WT1^+^/NPHS1^+^/CD31^+^) and CDH1^+^ tubular structures in day 18 organoids (**Figure 1B**) and compared morphology to mouse and human fetal kidney tissue. Day 11 organoids had cell clusters amongst mesenchymal tissue, and at day 14, discernable nephron-like structures (**Figure S1B**). At day 18, organoids had areas resembling the nephrogenic zone at embryonic day 19 (E19) in the mouse and between 8-10 wpc in human but lacked distinct cortico-medullary differentiation (**Figures 1C**, **S1A**-**B**). Using immunoelectron microscopy we confirmed the distribution of laminins in linear BM-like structures surrounding tubular segments in day 25 organoids and these were comparable to tubular BMs in the E19 mouse kidney (**Figure 1D**). Together, these findings demonstrate the presence of BM structures in kidney organoids.

**Figure 1.**
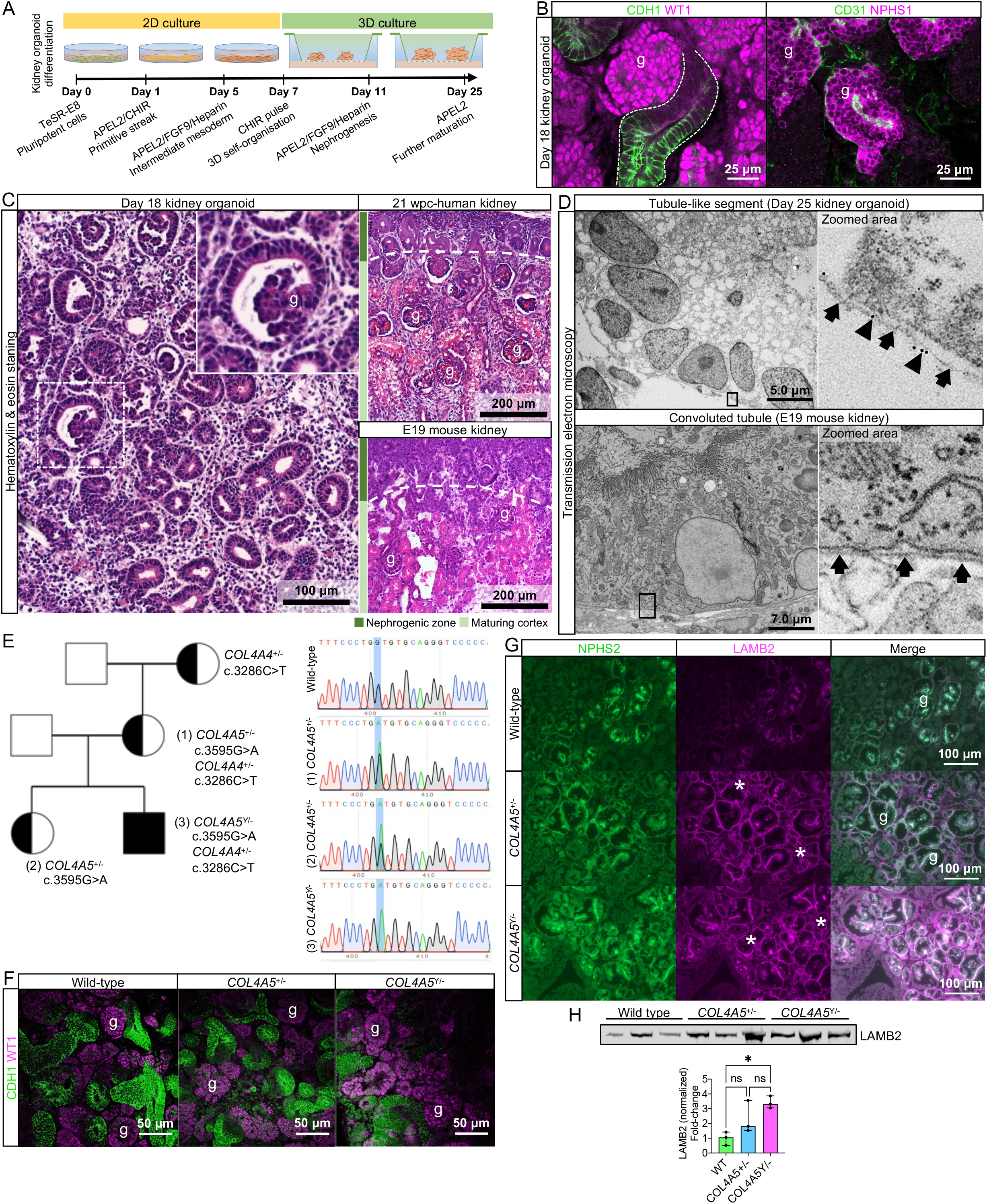
Kidney organoid basement membranes are altered in human disease. (A) Schematic representing the differentiation of iPSC’s to 3D kidney organoids. (B) Whole-mount immunofluorescence for kidney cell types: left image shows glomerular structures (g) with WT1^+^ cells and CDH1^+^ tubule segments (dashed line); right image shows a glomerular-like structure (g) containing podocytes (NPHS1^+^) and endothelial cells (CD31^+^). (C) Representative photomicrographs of day 18 kidney organoids (left) and human and mouse fetal kidneys (right) to demonstrate the comparable histological structure; inset shows an organoid glomerular structure (g). (D) Transmission electron photomicrographs of tubular BM in day 25 kidney organoid and E19 mouse fetal kidney: upper panels show a tubule-like structure in the organoid, and a BM-like sheet (arrows) labelled with a 10-nm gold-conjugated anti-laminin antibody (arrowhead); lower panel shows a murine convoluted tubule with a comparable tubular BM (arrows). (E) Pedigree from a family with missense variants in *COL4A5* and *COL4A4* genes. The sequencing data are shown for the *COL4A5* variant found in the mother and 2 siblings, which changes the amino acid from glycine to aspartic acid located in the triple-helical region of the collagen IV trimer. (F) Representative whole mount immunofluorescence images of wild-type and Alport kidney organoids show glomerular structures (g) containing WT1^+^ cells, and an intricate cluster of CDH1^+^ epithelial tubules. (G) Immunofluorescence for LAMB2 shows increased protein deposition in extraglomerular sites (*). NPHS2 was used as a podocyte marker to identify glomerular structures (g). (H) Immunoblotting for LAMB2 using total lysates from wild type and Alport organoids: bar chart shows relative fold change (to wild type). LAMB2 band optical density was normalized by Ponceau stain and compared by one-way ANOVA and Tukey’s multiple comparison tests (\**p*-value < 0.05; ns: not significant). Pooled data are presented as median, error bars indicate the 95% confidence interval for the median.

To determine the utility of organoids as a model to investigate abnormal BMs in kidney disease, we investigated iPSC lines from patients with Alport syndrome (AS), a genetic disorder caused by variants in type IV collagen genes (Barker et al., 1990). We selected iPSC lines from a mother and son, both carrying a pathogenic X-linked missense variant in *COL4A5* (c.3595G>A; p.Gly1232Asp) and a variant of unknown significance in *COL4A4* (c.3286C>T; p.Pro1096Ser; **Figure 1E**, see **Table S1** and **Supplementary information** for clinical details). AS patient-derived organoids progressed through differentiation and formed WT1^+^/NPHS1^+^/CDH1^+^ glomeruli and CDH1^+^ tubules, (**Figures 1F** and **S1C**) with no evident abnormalities by light microscopy. We found comparable distribution of collagen-α4 (IV) in AS and wild-type organoids (**Figure S1C**) confirming assembly of collagen IV α3α4α5, which is described in AS patients with missense variants (Yamamura et al., 2020a). Since laminin compensation is reported in X-linked AS (Abrahamson et al., 2007; Kashtan et al., 2001), we examined the deposition of laminin-β2 (LAMB2) in AS organoids. We found increased LAMB2 in AS organoids, most notable in extra-glomerular BM (**Figure 1G**), and further confirmed increased LAMB2 levels with immunoblotting (**Figure 1H**). Together, these findings demonstrate the utility of kidney organoids to reveal abnormal patterns of BM assembly in human development and disease.

### A conserved sequence of BM assembly in kidney organoids

Having identified BM structures in kidney organoids, we next explored the potential for this system to model human BM assembly. Studies in mouse and invertebrate development have shown a sequence of BM assembly with initial laminin deposition followed by incorporation of collagen IV, nidogen and perlecan (Jayadev et al., 2019; Matsubayashi et al., 2017; Urbano et al., 2009). To investigate the assembly sequence in organoids, we used whole-mount immunofluorescence to examine the temporal co-deposition of COL4A1 with laminin (using a pan-laminin antibody), and nidogen with perlecan during differentiation. We found punctate deposits of pan-laminin in interrupted BM networks around cell clusters in day 11 organoids, and in continuous BMs around CD31^+^ endothelial and epithelial structures in day 18 and 25 organoids (**Figure 2A**). Conversely, COL4A1 was weakly detected in day 11 organoids, partially co-distributed with laminin by day 18, and within continuous BM networks by day 25 (**Figure 2A**). Nidogen and perlecan colocalized in discreet, interrupted BMs in day 11 organoids and later in linear BM networks around tubules and NPHS1^+^ glomerular structures on day 18 and 25 (**Figure 2B**). Together, these findings indicate that kidney organoids recapitulate the sequence of BM assembly described *in vivo*, reinforcing their fidelity as system for investigating BM dynamics.

**Figure 2.**
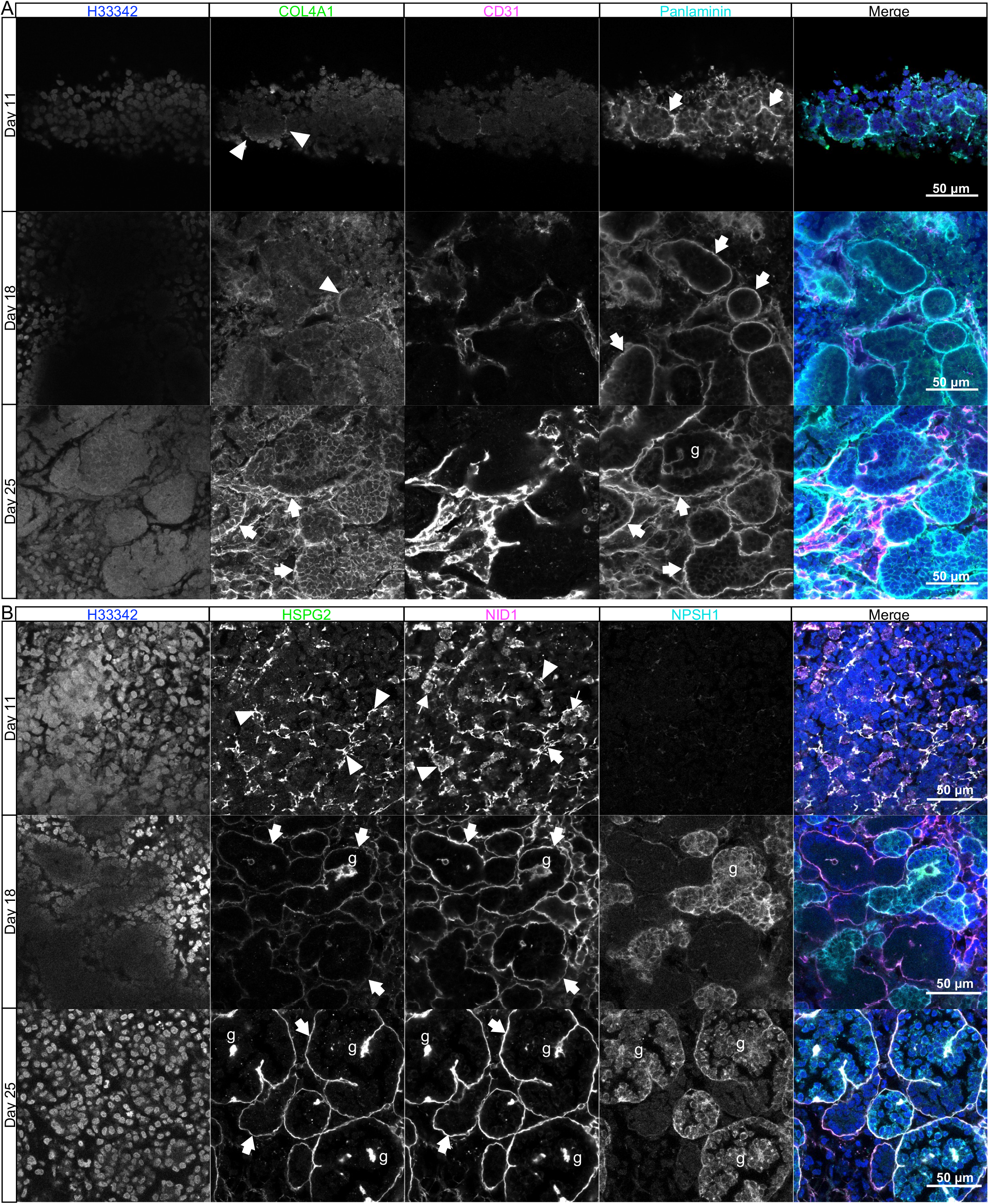
Sequential assembly of basement membranes components. (A) Confocal immunofluorescence microscopy of wild-type kidney organoids showing the temporal emergence and BM co-distribution of COL4A1 and panlaminin and (B) perlecan and nidogen on days 11, 18 and 25 of differentiation. NPHS1 and CD31 were used respectively as markers for podocyte and endothelial cells in glomerular like structures (g). Arrowheads indicate interrupted BM segments, large arrows indicate diffuse BM networks, and thin arrows indicate intracellular droplets of BM proteins.

### Time course proteomics reveals complex dynamics of BM assembly

To understand global BM dynamics, we investigated organoids at day 14, 18 and 25 with time course proteomic analysis. We broadly separated intracellular and extracellular proteins by fractionation (**Figures 3A, 3B and S2A**) based on solubility (Lennon et al., 2014). Overall, we detected 5,245 proteins in the cellular fraction and 4,703 in the extracellular fraction (**Table S2**), and by cross-referencing with the human matrisome (Naba et al., 2016) we identified 228 matrix proteins in kidney organoids (**Figure S2B**, **Table S2**). Principal component analysis highlighted discreet clustering for the organoid time points based on matrix protein abundance (**Figure S2C**). There was an increase in matrix abundance from day 14 to 25 (**Figure 3B**) and 203 (~90%) of matrix proteins were detectable at all time points (**Figure 3C**). This initial analysis confirmed a gradual assembly of matrix during organoid differentiation. To address global BM composition, to identified BM proteins using the comprehensive BM gene network curated in *basement membrane*BASE (https://bmbasedb.manchester.ac.uk/, updated in 2021). The organoid extracellular fraction was enriched for BM proteins compared to the cellular fraction (**Figure 3D**), which was expected as these are large, highly cross-linked proteins and hence, difficult to solubilize. Furthermore, we observed an increasing trend for BM protein levels through day 14 to 25 (**Figure 3D**), again indicating BM deposition over time, and corroborating our previous immunofluorescence findings. In total, we identified 78 BM proteins (**Figure 3E**) including components deposited early in kidney morphogenesis (*e.g.*, COL4A1, COL4A2, LAMA1, LAMB1, LAMC1) (**Figures 3F** and **S2D**). LAMA5 and LAMB2, two key components of the mature GBM, only appeared amongst the most abundant BM components at day 25 (**Figure S2D**) indicating a temporal expression of GBM laminins during organoid differentiation (**Figure 3F**). This was confirmed by marked upregulation of mature GBM proteins from day 18 to 25, with LAMB2 scoring with the highest fold-change followed by LAMA5 and COL4A3 (**Figures 3G and 3H**). LAMA5 was also enriched from day 14 to 18 together with other GBM proteins (COL4A3, AGRN) and early BM collagens and laminins (COL4A1, COL4A2, LAMC1, LAMA1) (**Figures 3G, 3H and S2E**).

**Figure 3.**
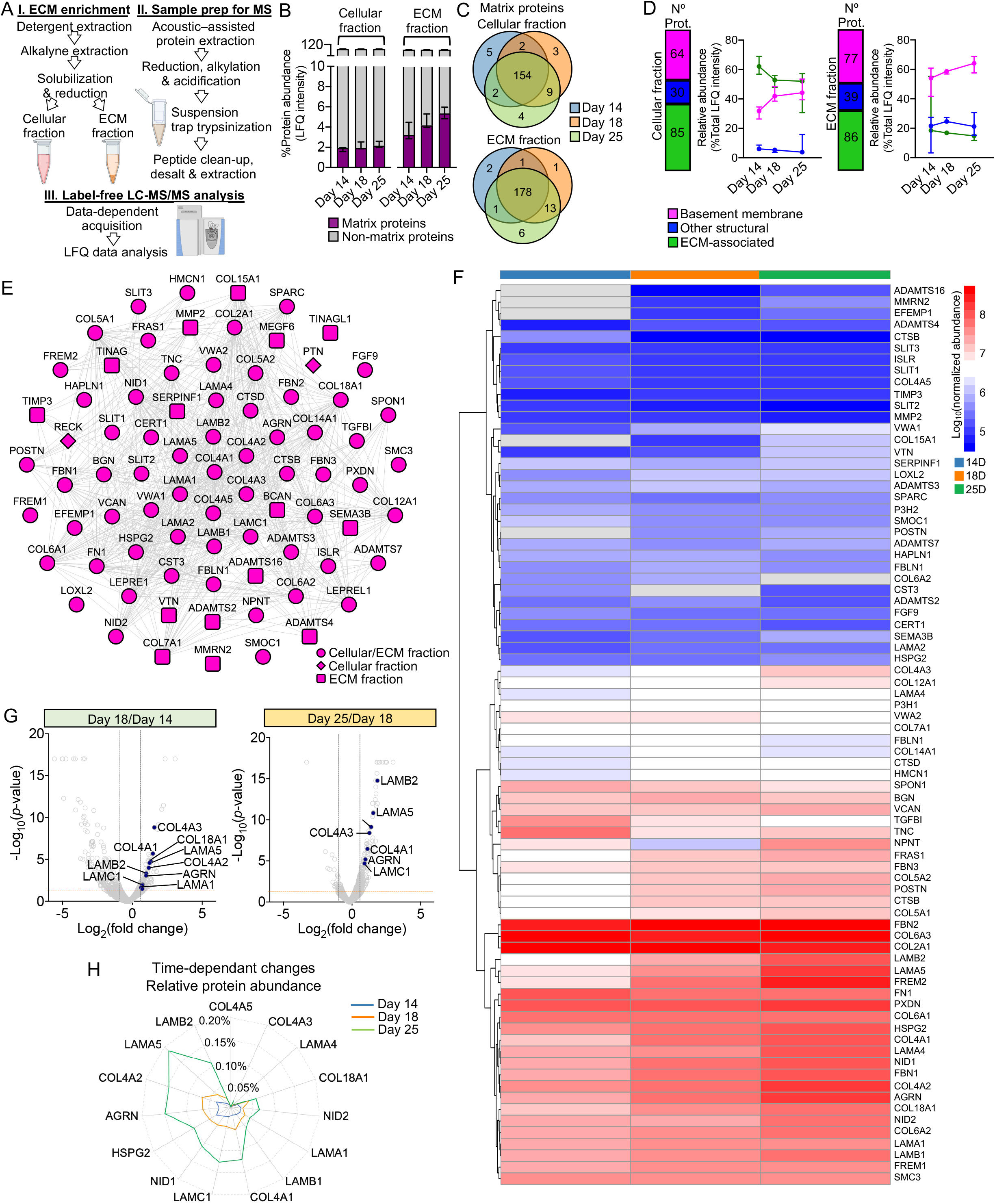
Time course proteomics reveals complex dynamics of basement membrane assembly. (A) Schematic for sample enrichment for matrix (ECM) proteins for tandem MS analysis (created with BioRender.com). (B) Bar graphs show the relative abundance of matrix proteins and non-matrix proteins identified by MS analysis in the cellular and ECM fractions of kidney organoids on days 14, 18 and 25. Pooled data are presented as median, error bars indicate the 95% confidence interval for the median. (C) Venn diagrams showing identification overlap for matrix proteins detected in organoids on days 14, 18 and 25. (D) Matrix proteins are classified as basement membrane, other structural and ECM-associated proteins. Bar charts show the number of matrix proteins per each category in both cellular and ECM fractions, and line charts show the changes in the relative abundance (percentage of total matrix abundance) for the matrix categories over the time course differentiation. Pooled data are presented as median, error bars indicate the 95% confidence interval for the median. (E) Protein interaction network showing all BM proteins identified over the kidney organoid time course MS study (nodes represent proteins and connecting lines indicate protein-protein interaction). (F) Heat map showing the log_10_-transformed abundance levels of BM proteins identified in the ECM fraction along kidney organoid differentiation time course (proteins detected only in one time-point are not shown). (G) Volcano plots show the log_2_-fold change (x-axis) versus −log_10_-*p*-value (y-axis) for proteins differentially expressed in the ECM fraction of kidney organoids from day 14 to 18, and 18 to 25. Key BM proteins significantly up-regulated (FC ≥ 1.5, *p*-value < 0.05, Two-way ANOVA test, *n*=3) are indicated. (H) Time-dependant changes in the relative abundance (percentage of total protein intensity) of key BM proteins in the ECM fraction of kidney organoids during differentiation. Pooled data are presented as median.

During GBM assembly, an initial laminin-111 (α1αβ1γ1) network is sequentially replaced by laminin-511 then −521 (Abrahamson et al., 2013). We therefore reasoned that day 14 to 18 would represent a period of intense BM assembly and initial GBM differentiation. In support of this hypothesis, a pathway enrichment analysis of upregulated proteins from day 14 to 18 revealed an overrepresentation of terms associated with BM assembly and remodeling, including laminin interactions, degradation of ECM and collagen chain trimerization (**Figure S2F**). Together, this global proteomic analysis revealed new insights into the complexities of BM dynamics and the distinct temporal emergence of BM isoforms required for long term functional integrity.

### Tracking collagen IV and laminin isoforms during organoid differentiation

To confirm the temporal emergence of specific BM isoforms we investigated the distribution of COL4A1, COL4A3, LAMA5, LAMB1, LAMB2 in organoid BMs through day 14 to 25 by immunofluorescence (**Figure 4D**). As described earlier, COL4A1 appeared on day 11 (**Figure 2A**) and partial colocalization with laminin on day 14, and as continuous BM networks from day 18 (**Figure 4A**). Conversely, COL4A3 was scarce from day 14 to 18, but clearly colocalized with laminin in glomerular structures on day 25 (**Figure 4A**). We detected LAMA5 from day 18 and this increased in glomerular structures at day 25. LAMB1 was widely distributed from day 14 to 25 whereas LAMB2, detected from day 18 onwards, was enriched in glomerular structures at day 25. These findings not only confirm a temporal emergence of specific BM isoforms, but also highlight specific localization to glomerular structures later in differentiation.

**Figure 4.**
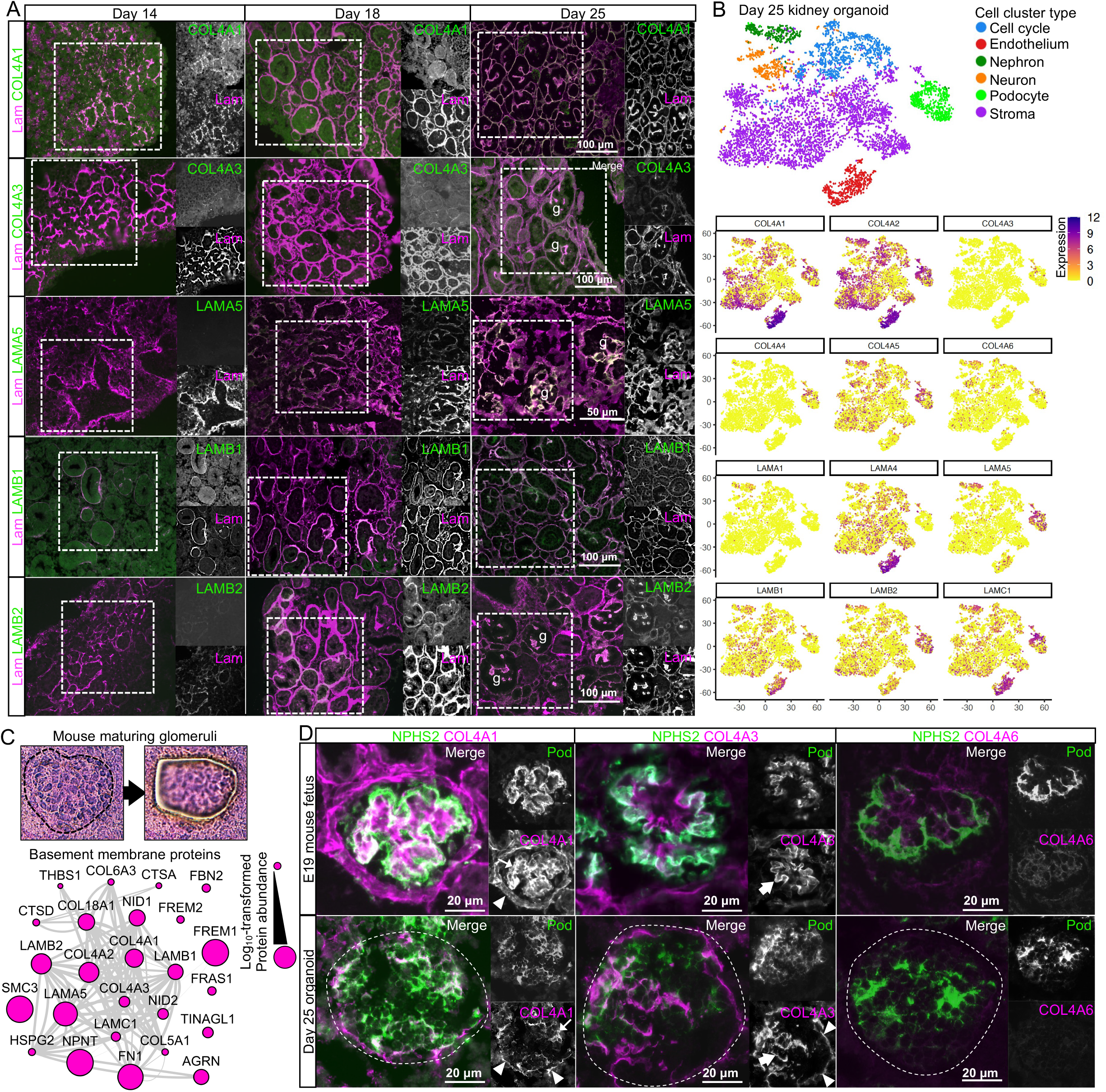
Key collagen IV and laminin isoform transitions occur during kidney organoid differentiation. (A) Immunofluorescence for key type IV collagen and laminin isoforms showing their emergence and distribution in kidney organoid BM. Panlaminin antibody was used to co-label organoid BM; glomerular structures are indicated (g). (B) Re-analysis of a kidney organoid scRNA-seq dataset (GSE114802, (Combes et al., 2019a) confirms cellular specificity for collagen IV and laminin isoform gene expression. tSNE plots represent the cell type clusters identified, and colour intensity indicate the cell-specific level of expression for the selected BM genes. (C) Proteomic analysis of laser-captured maturing glomeruli from E19 mouse kidneys. Histological images show the laser dissected glomeruli, and the protein interaction network shows 25 BM proteins identified (nodes represent proteins and connecting lines indicate protein-protein interaction). (D) Immunofluorescence for specific collagen IV isoforms in maturing glomeruli in E19 mouse kidney and in glomerular structures (indicated by dashed lines) in day 25 organoids. NPHS2 was used to label podocytes. Arrowheads indicate outer Bowman’s capsule (mouse) or glomerular surface (organoid), large arrows indicate GBM (mouse) or GBM-like deposition (organoid); thin arrows indicate mesangial matrix (mouse) or internal glomerular deposition (organoid).

We then hypothesized that distinct cell types would express specific BM isoforms to concentrate their distribution and therefore analyzed single-cell RNA sequencing (scRNA-seq) data from day 25 kidney organoids (Combes et al., 2019a) to map the expression profile for BM genes (**Figure S3A**, **Table S3**). We found NPHS2^+^/PODXL^+^ podocytes were the main source of COL4A3, COL4A4 (**Figures 4B and S3A**) and also had high levels of expression for LAMA5 and LAMB2. PECAM1^+^/KDR^+^ endothelial cells, MAB21L2^+^/CXCL14^+^ stromal cells and PAX8^+^/PAX2^+^ nephron cell lineages all expressed COL4A1, COL4A2, LAMB1, LAMC1 and LAMA1 was detected in the nephron cell cluster whereas LAMA4 was expressed by both endothelial and stroma cells (**Figures 4B and S3A**). These findings align with current understanding of kidney development *in vivo* and indicate that kidney organoids recapitulate the known cell specific contributions to BM assembly during glomerulogenesis.

Since the developmental transition from the α1α1α2 to the α3α4α5 network of collagen IV is key for long term GBM function and reduced or absent α3α4α5 leads to loss of function (Miner and Sanes, 1996), we mapped the localization of collagen IV isoforms in day 25 organoids and compared to E19 mouse glomeruli. We performed laser microdissection and proteomic analysis of maturing mouse glomeruli (**Table S4**) and identified 25 BM proteins (also detected in kidney organoids), including COL4A1, COL4A2 and COL4A3 (**Figure 4C**) thus indicating presence of α1α1α2 and α3α4α5 networks. We compared the localization of COL4A1 and COL4A3 by immunofluorescence in the mouse and NPHS2^+^ glomeruli in kidney organoids (**Figure 4D**). We observed a similar distribution of COL4A1, but for COL4A3, we found GBM-like and extraglomerular distribution in kidney organoids (**Figure 4D**). Therefore, kidney organoids initiate collagen IV isoform transitions during glomerulogenesis. To further explore mechanisms of transition, we investigated the expression of LIM homeodomain transcription factor 1-beta (LMX1b) and FERM-domain protein EPB41L5 in day 25 organoids (**Figure S3B**). LMX1b and EPB41L5 are proposed regulators of GBM assembly and isoform transitions during development (Maier et al., 2021; Morello et al., 2001). Moreover, EPB41L5 is also implicated in regulating the incorporation of laminin-511 and −521, into stable GBM scaffolds. We found that similar cell populations expressed EPB41L5 and LAMA5 (**Figure S3B**), and there was an increase in EPB41L5 protein levels from day 14 to 18 (**Figure S3C**), coinciding with an increase in LAMA5 (**Figures 3G-H**). Together, these findings demonstrate that kidney organoids initiate isoform transitions during glomerular differentiation, with the expression of known BM regulators.

### BMs in late-stage organoids and fetal kidneys are highly correlated

To relate BM assembly in organoids to a comparable *in vivo* system, we examined fetal kidneys. Having verified morphological similarity between day 25 organoids and E19 mouse fetal kidney (**Figure 1C**), we used this timepoint for comparison by proteomic analysis. We generated cellular and extracellular fractions from whole fetal kidneys (**Figures 5A**, **S4A** and **S4B**) and identified 208 matrix components from a total of 5,071 proteins (**Figure 5A**; **Table S4**). These included 83 BM proteins and the most abundant were those seen early (COL4A1, COL4A2, COL18A1, LAMB1, LAMC1) and later (LAMA5) in BM assembly (**Figure S5C**). We compared these findings to a proteomic dataset from the E18.5 mouse kidney (Lipp et al., 2021), and found considerable overlap, with a further 130 matrix protein identifications (**Figure 5C**), including key GBM components COL4A3, COL4A4, COL4A5 and COL4A6. We then compared the BM proteins from each organoid timepoint with E19 mouse kidneys (**Figure 5D**) and found the highest overlap (58.4%) between E19 and day 25 kidney organoids. In line with previous findings (Hale et al., 2018), this comparison highlighted later expression of TINAG and TINAGL1 in day 25 organoids, also detected in E19 mouse kidneys but not in day 14 or day 18 organoids. TINAGL1 was also detected in E19 mature glomeruli (**Figure 4C**) and in proteomic studies of adult human glomeruli (Lennon et al., 2014) but its role for BM biology remains unknown. In addition, the overlap for other structural and matrix-associated proteins was lower than for BM proteins (**Figure S4D**), which highlights conservation of BM composition between mouse and kidney organoids. To further verify similarities, we performed a Spearman’s rank correlation and found E19 had the higher correlation with more differentiated organoids (**Figures 5E and S4E**).

**Figure 5.**
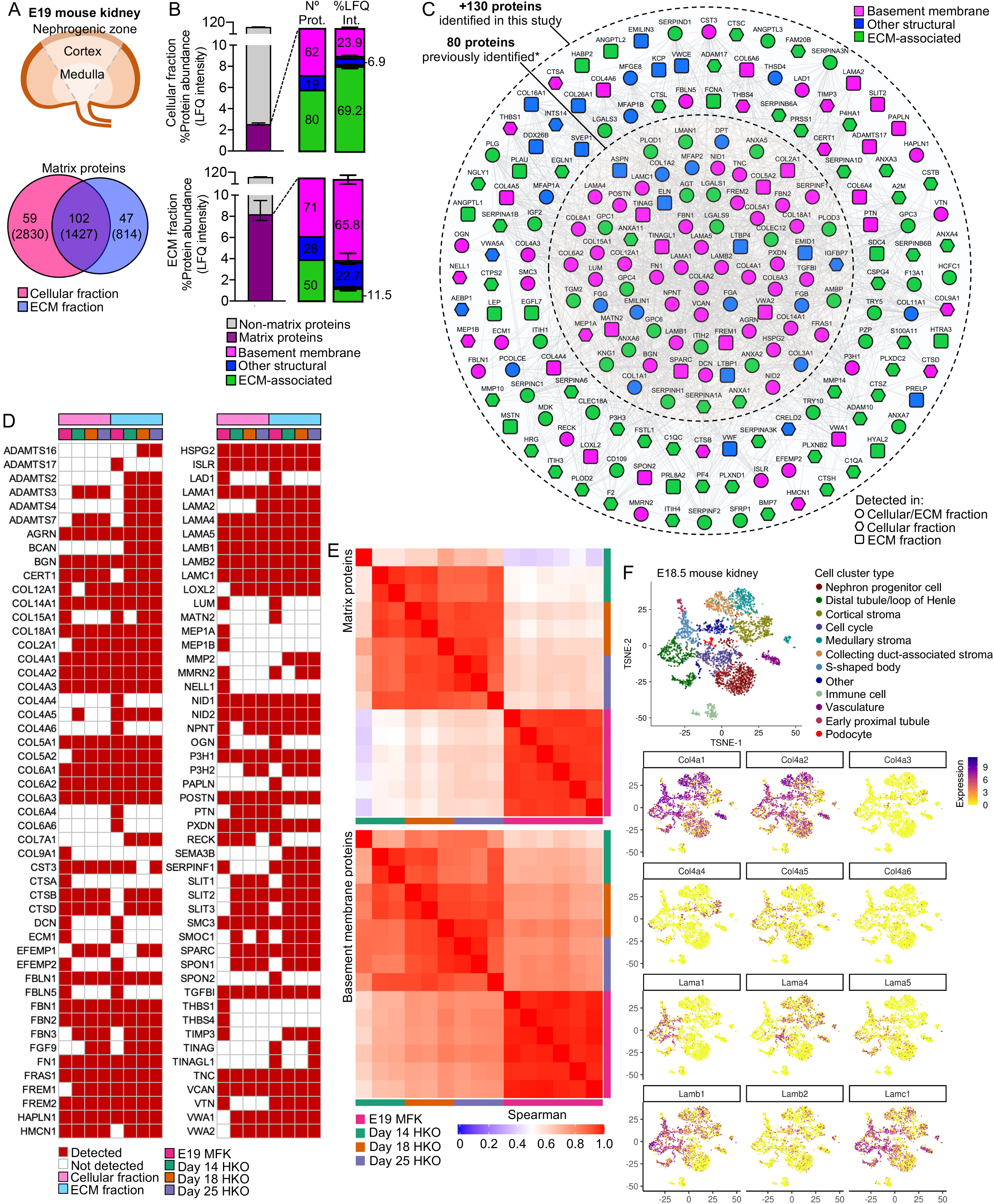
Basement membranes in mouse fetal kidneys are comparable to kidney organoids. (A) Schematic representation of the E19 mouse kidney sampled for MS-based proteomics, and a Venn diagram showing the overlap for matrix proteins identified in the cellular and ECM fractions. (B) Bar charts show enrichment for matrix proteins in both cellular and ECM fractions, as indicated by the number and relative abundance of proteins in each matrix category. Pooled data are presented as median, error bars indicate the 95% confidence interval for the median. (C) Expanded fetal mouse kidney matrisome represented as a protein interaction network (nodes represent proteins identified in this and in a previous study (Lipp et al., 2021), and connecting lines indicate protein-protein interaction). (D) Comparison of BM proteins identified in the E19 mouse fetal kidney* (MFK) and human kidney organoids (HKO) during differentiation (*corresponding human ortholog proteins are shown). (E) Spearman rank correlation analysis for matrix and BM protein abundance (ECM fraction) comparisons between E19 mouse fetal kidney (MFK) and human kidney organoids (HKO). (F) Re-analysis of a E18.5 mouse kidney scRNA-seq dataset (GSE108291, (Combes et al., 2019b) confirms cellular specificity for collagen IV and laminin isoform gene expression. tSNE plots represent the cell type clusters identified, and colour intensity indicate the cell-specific level of expression for the selected BM genes.

Next we analyzed scRNA-seq datasets from E18.5 mouse kidneys ((Combes et al., 2019b); **Figure 5F**) and from 8- and 9-wpc human kidneys ((Young et al., 2018); **Figure S5**) to identify cells expressing specific BM genes (**Table S3**). In the mouse we found mature GBM components expressed by Synpo^+^/Nphs2^+^ podocytes (Col4a3, Col4a4, Lama5, Lamb2), Plvap^+^/Pecam1^+^ vascular cells (Lama5, Lamb2), and Cited1^+^/Crym^+^ nephron progenitor cell lineages, and Aldob^+^/Fxyd2^+^ tubular, vascular and Six2^+^ stromal cells predominantly contributing with Col4a1, Col4a2, Lamb1 and Lamc1 expression (**Figure 5F**). Lama1 was mainly expressed by Clu^+^/Osr2^+^ S-shaped bodies and Gata3^+^/Wfdc2^+^ ureteric bud/distal tubular cells, and Lama4 restricted to vascular and stromal cells. A similar pattern was observed in the embryonic human kidney (**Figure S5**), and these findings were also consistent with our findings in day 25 kidney organoids (**Figure 4B** and **S3**). Interestingly, immune cells also contributed for Col4a1 and Col4a2 expression in both mouse and human kidneys. Collectively, these findings highlight the conservation of BM gene expression across organoids and fetal kidneys.

### Basement membranes are dynamic through embryonic development to adulthood

Having observed dynamic BM composition during kidney development we then compared composition in adulthood. For this, we analyzed proteomic data from isolated adult human nephron compartments and identified 71 glomerular BM proteins, and 61 in the tubulointerstitium (**Table S6**). We compared BM networks in the adult kidney with both developmental systems (day 25 kidney organoids and E19 mouse kidney) and found a significant overlap with 44 of 107 BM proteins shared amongst all datasets. These included core components (COL4A1, COL4A2, LAMA1, LAMB1, LAMC1, HSPG2, COL18A1, NID1, NID2), mature GBM components (COL4A3, COL4A5, LAMA5, LAMB2) and many minor structural proteins (**Figure 6A**). We found a strong correlation between both matrix and BM networks in glomerular and tubulointerstitial compartments, but lower correlation between adult and developmental datasets (**Figures 6B** and **S6A**). Interestingly, the correlation between adult and development data was lower for BM than for all matrix proteins, suggesting that although kidney BMs retain a consistent profile through development to adulthood, there is diversification within distinct kidney compartments.

**Figure 6.**
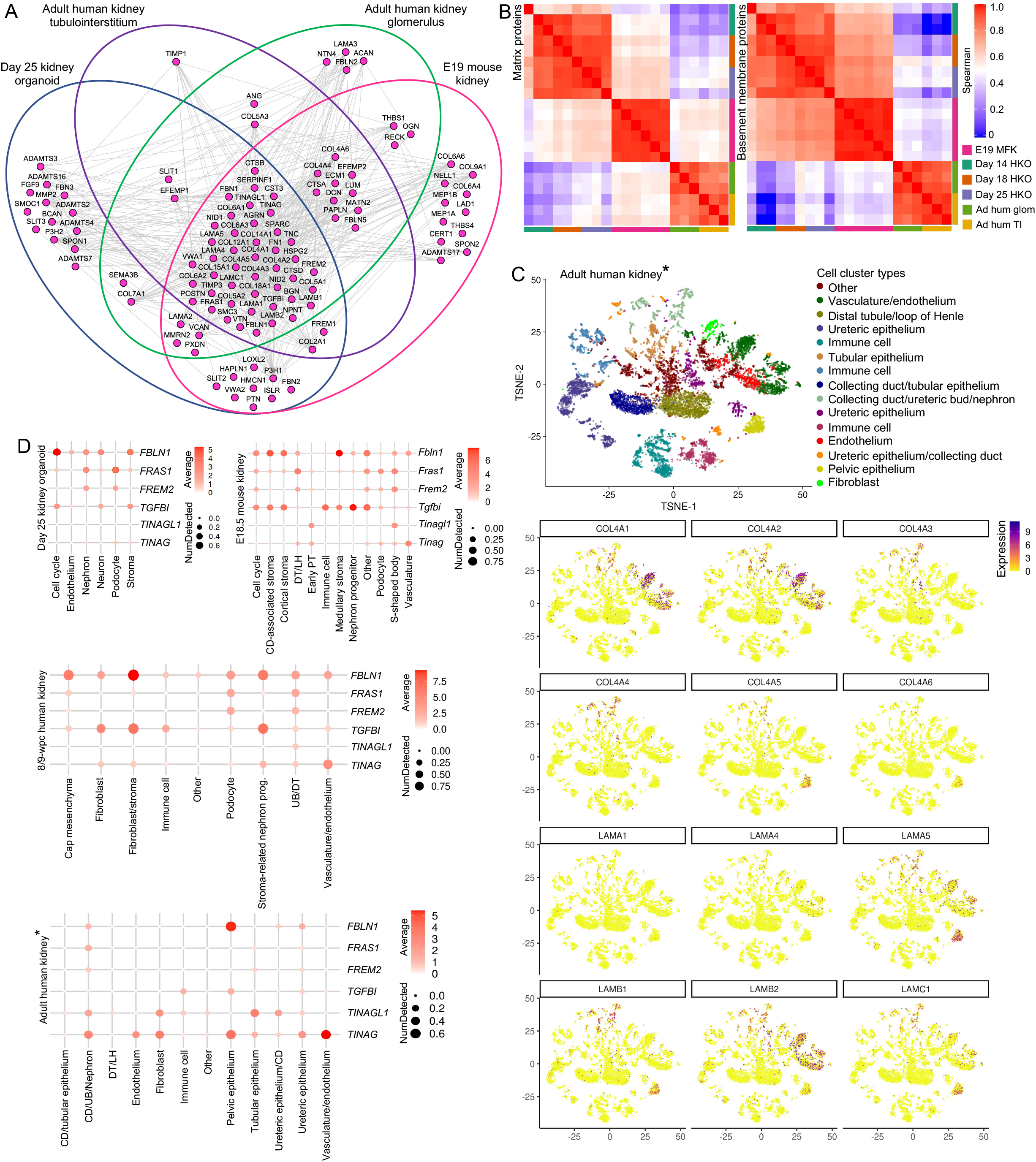
Basement membranes are dynamic through embryonic development to adulthood. (A) Integrative interactome showing a common core of 44 BM proteins across day 25 organoid, E19 mouse kidney, adult human glomerular and tubulointerstitial compartments. Venn diagrams indicate in which dataset each BM protein was detected. Nodes represent proteins, and lines indicate protein-protein interactions. (B) Spearman rank correlation analysis for matrix and BM protein abundance (ECM fraction) comparisons between E19 mouse fetal kidney (MFK), human kidney organoids (HKO) and adult human glomerulus and kidney tubulointerstitium. (C) Re-analysis of an adult human kidney scRNA-seq dataset (EGAS00001002553, (Young et al., 2018)) confirms cellular specificity for collagen IV and laminin isoform gene expression. tSNE plots represent the cell type clusters identified, and colour intensity indicate the cell-specific level of expression for the selected BM genes (*proximal tubule cells were not included). (D) Cell expression of minor BM components through kidney development to adulthood. Dot plots show the level of expression of target genes in all published datasets re-analysed in this study (*proximal tubule cells were not included).

To understand cellular origins of BM components through development to adulthood, we analyzed adult human kidney scRNA-seq data (Young et al. 2018; **Table S3**), and as with organoids and mouse fetal kidney, we found COL4A1, COL4A2, LAMB1 and LAMC1 predominantly expressed by endothelial and tubular cells. Although podocyte markers were not enriched in the adult dataset, we detected COL4A3 and COL4A4 expression by PAX8 ^+^ nephron cell types, and LAMA5 and LAMB2 mainly expressed by KRT5^+^/EMCN^+^ endothelial cells (**Figure 6C**). LAMA4 was widely detected in both E18.5 mouse kidney and day 25 organoids (**Figures 3B, 4F** and **S3**), but barely present in the adult kidney (**Figure 6C**), consistent with previous reports of transient expression in human kidney development (Miner, 1999). These findings demonstrate consistent cellular origins for BM components through development to adulthood.

To further verify the extent of this consistency, we selected minor BM components across all scRNA-seq datasets (**Figure 6D**). These included: FBLN1, TGFBI both implicated in BM remodeling (Boutboul et al., 2006; Feitosa et al., 2012); FRAS1, FREM2 important for branching morphogenesis (Chiotaki et al., 2007; Petrou et al., 2008); and TINAG, TINAGL1, with unknown roles in BM function. We found a common pattern of enrichment amongst the developmental datasets for FBLN1, which was expressed by stromal cells; FRAS1 and FREM2 were expressed by podocyte and nephron cell clusters; and TGFBI by stromal cells, nephron progenitors, and also by immune cells. In human adult kidneys FBLN1 was mainly present in pelvic epithelial cells and FRAS1/FREM2 in ureteric bud/distal tubule cells, thus indicating spatiotemporal expression of these components. TINAG and TINGAL1 had variable patterns of cell expression across datasets. This comparative analysis shows conservation of distinct cell types contributing to BM assembly during kidney development and uncovers diversification in cellular origins in adult kidneys.

## DISCUSSION

BMs are complex structures and proteomic studies have highlighted this complexity in homeostasis and disease (Lennon et al., 2014; Randles et al., 2015). During development, BMs are also highly dynamic and undergo intense remodeling (Kyprianou et al., 2020) but understanding of BM assembly and regulation is limited by lack of appropriate systems to track components that are spatiotemporally regulated and undergoing turnover (Naylor et al., 2020). Human fetal tissue has limited availability and studies are restricted to static time points and the technical limitations of imaging in mouse models also impairs investigation of the dynamic BM environment *in vivo*. However, BM studies in *Drosophila* and *C. elegans* have provided important insights into BM dynamics and turnover using fluorescent tagging of endogenous proteins (Keeley et al., 2020; Matsubayashi et al., 2020) and highlight the power of studying BM dynamics. Here we demonstrate the fidelity of kidney organoids, making it an exceptional system for investigating BM assembly and regulation in health and disease.

Despite the developmental limitations with iPSC-derived kidney organoids, including lack of directional cues and cortical-medullary segmentation (Romero-Guevara et al., 2020), this system has morphological and molecular features comparable to fetal kidney tissue that overcome species differences and provides a complex *in vitro* environment to examine BM assembly (Bantounas et al., 2018). We demonstrated that kidney organoids differentiate into glomerular structures containing the cells required for GBM assembly. Single-cell transcriptomic studies have also shown over 20 other distinct cell populations in kidney organoids (Combes et al., 2019b; Wu et al., 2018). Cross-talk between different cell types is essential for BM assembly and influences composition (Byron et al., 2014) hence, the multiple cell types in the organoid system enable BM formation and remodeling. We discovered that organoids form BMs early during differentiation, and more importantly, recapitulate a sequence of assembly events with initial deposition of laminin followed by incorporation of type IV collagen, nidogen and perlecan (Brown et al., 2017; Sasaki et al., 2004).

Kidney organoids have also provided new insights into developmental programmes and disease processes (Rooney et al., 2021; Tanigawa et al., 2018; Tian and Lennon, 2019). We found that organoids with a pathological missense variant in *COL4A5* differentiated and deposited core BM proteins, including a collagen α3α4α5 network, which is described in missense variants. In one study 64 out of 146 patients with X-linked Alport syndrome had collagen-α5 (IV) in the GBM (Yamamura et al., 2020b). Despite evidence of protein secretion, the GBM fails to maintain function in these patients. Interestingly, we found increased deposition of LAMB2 in extraglomerular BMs in Alport organoids. Dysregulation of glomerular laminins, including LAMB2, in patients with Alport syndrome and animal models has been reported (Abrahamson et al., 2007; Kashtan et al., 2001), but the mechanisms are unclear. Our findings in Alport organoids demonstrate the utility of this system to dissect abnormal mechanisms of BM assembly.

To define global BM composition in kidney organoids we used MS-based proteomics and identified 202 BM proteins dynamically expressed throughout the differentiation time course. Core GBM components including laminin-521 and collagen α3α4α5(IV), which only appear in mature glomeruli, were also detected. Developmental isoform transitions in the GBM involving laminin and type IV collagen are described in humans and rodents (Abrahamson and St John, 1993) and in current understanding, immature glomeruli assemble a primary GBM containing laminin-111 and collagen α1α2α1(IV) that is later replaced by laminin-521 and collagen α3α4α5(IV) in mature glomeruli. In keeping with these observations, we found a temporal and spatial emergence of mature collagen IV and laminin isoforms within glomerular structures in kidney organoids, and in scRNA-seq data we confirmed podocyte expression of mature GBM markers. These findings demonstrate that kidney organoids can efficiently recapitulate the spatiotemporal emergence of GBM components. The triggers for isoform switching remains unknown but two regulators, LMX1b and EPB41L5, have been implicated. Studies in *Lmxb1* knockout mice suggest that podocyte expression of this transcription factor is not essential for initial GBM assembly but linked to *Col4a3*/*Col4a4* expression during glomerulogenesis as demonstrated by reduced collagen α3α4α5(IV) network in the GBM in null *Lmxb1* newborn mice (Morello et al., 2001). We found podocyte specific expression of *LMX1b*/*COL4A3*/*COL4A4* in day 25 kidney organoids, and moreover, confirmed deposition of COL4A3/COL4A4 in a GBM pattern within NPHS2^+^-glomerular structures in organoids. In addition, podocyte expression of EPB41L5, a component of the podocyte integrin adhesion complex, was linked to GBM assembly *in vivo* and incorporation of laminin-511/521 into extracellular BM networks *in vitro* (Maier et al., 2021). In this study, we detected EPB41L5 in organoid and mouse proteomic datasets, with cell expression patterns matching LAMA5, indicating the potential for this system to unravel further insights into GBM regulation.

There are few proteomic studies addressing the spatiotemporal changes in BM during development. One study of mouse kidney development, defined the global ECM composition through development to adulthood and described a sequence of changes in interstitial matrix over development (Lipp et al., 2021). With our sample fractionation strategy and analytical pipeline, we detected a further 130 matrix proteins, including key GBM components (COL4A3, COL4A4, COL4A5, COL4A6) and BM regulators such as PAPLN and HMCN1 (Keeley et al., 2020; Morrissey et al., 2014). Our findings further demonstrated that late-stage kidney organoids (at day 25) and mouse kidneys on E19 share a very comparable BM profile. We also verified similar patterns of BM gene expression between kidney organoids and fetal human and mouse kidneys, especially for type IV collagen and laminin isoforms. Overall these data demonstrate the high-fidelity with which kidney organoids recapitulate BM gene expression and protein composition seen *in vivo*. Thus, we conclude that kidney organoids are a highly tractable model that can be used to study the dynamic nature of human BM assembly in both health and disease.

## ACKNOWLEDGMENTS

This work was supported by a Wellcome Senior Fellowship award (202860/Z/16/Z) to R.L., a Kidney Research UK grant (RP52/2014) awarded to R.L to support a PDRA position for P.T. M.R.P.T.M. was supported by the São Paulo Research Foundation (FAPESP, #2015/02535-2; #2017/26785-5) and by the Global Challenge Research Fellowship program. R.Z. is supported by Veterans Affairs Merit Awards 1I01BX002196-01 and DK069221. DAL’s laboratory is supported by the NIHR Biomedical Research Centre at Great Ormond Street Hospital for Children NHS Foundation Trust and University College London, an Investigator Award from the Wellcome Trust (220895/Z/20/Z) and project grants from the Medical Research Council (MR/P018629/1 and MR/J003638/1). The mass spectrometer and microscopes used in this study were purchased with grants from the Biotechnology and Biological Sciences Research Council, Wellcome Trust and the University of Manchester Strategic Fund. Mass spectrometry was performed in the Biomolecular Analysis Core Facility, Faculty of Life Sciences, University of Manchester, and we thank David Knight, Ronan O’cualain and Stacey Warwood for advice and technical support and Julian Selley for bioinformatic support. The iPSC lines were generated at the Wellcome Trust Sanger Institute, under the Human Induced Pluripotent Stem Cell Initiative (HipSci) funded by a grant (WT098503) from the Wellcome Trust and Medical Research Council. We also acknowledge Faris Tengku who helped with the generation of wild type iPSC and Joseph Luckman who helped to optimise immunofluorescence protocols and Karen Leigh Price and Maria Kolatsi-Joannou who helped with the human histology.

## AUTHORS CONTRIBUTIONS

Study conceptualization and experimental design: MRPTM, PT, RL; Experimental procedures, data acquisition and analysis: PT, MRPTM, CL, AM, SW; Manuscript preparation: MRPTM, PT, LH, DG, RL; Manuscript editing and review: MRPTM, PT, LH, SW, DL, DG, SK, RZ, RL; single cell RNA sequencing analyses: SMB, CL; Provided patient samples for iPSC: DG; Provided fetal human kidney sections: DAL; Provided fetal mouse kidney samples: MRPTM, TMTZ.

## References

Abrahamson, D.R., and St John, P.L. (1993). Laminin distribution in developing glomerular basement membranes. Kidney Int. 43, 73–78.

Abrahamson, D.R., Isom, K., Roach, E., Stroganova, L., Zelenchuk, A., Miner, J.H., and St John, P.L. (2007). Laminin compensation in collagen alpha3(IV) knockout (Alport) glomeruli contributes to permeability defects. J. Am. Soc. Nephrol. 18, 2465–2472.

Abrahamson, D.R., St John, P.L., Stroganova, L., Zelenchuk, A., and Steenhard, B.M. (2013). Laminin and type IV collagen isoform substitutions occur in temporally and spatially distinct patterns in developing kidney glomerular basement membranes. J. Histochem. Cytochem. 61, 706–718.

Bantounas, I., Ranjzad, P., Tengku, F., Silajdžić, E., Forster, D., Asselin, M.-C., Lewis, P., Lennon, R., Plagge, A., Wang, Q., et al. (2018). Generation of Functioning Nephrons by Implanting Human Pluripotent Stem Cell-Derived Kidney Progenitors. Stem Cell Rep. 10, 766–779.

Barker, D.F., Hostikka, S.L., Zhou, J., Chow, L.T., Oliphant, A.R., Gerken, S.C., Gregory, M.C., Skolnick, M.H., Atkin, C.L., and Tryggvason, K. (1990). Identification of mutations in the COL4A5 collagen gene in Alport syndrome. Science 248, 1224–1227.

Bonnans, C., Chou, J., and Werb, Z. (2014). Remodelling the extracellular matrix in development and disease. Nat. Rev. Mol. Cell Biol. 15, 786–801.

Boutboul, S., Black, G.C.M., Moore, J.E., Sinton, J., Menasche, M., Munier, F.L., Laroche, L., Abitbol, M., and Schorderet, D.F. (2006). A subset of patients with epithelial basement membrane corneal dystrophy have mutations in TGFBI/BIGH3. Hum. Mutat. 27, 553–557.

Brown, K.L., Cummings, C.F., Vanacore, R.M., and Hudson, B.G. (2017). Building collagen IV smart scaffolds on the outside of cells. Protein Sci. 26, 2151–2161.

Byron, A., Randles, M.J., Humphries, J.D., Mironov, A., Hamidi, H., Harris, S., Mathieson, P.W., Saleem, M.A., Satchell, S.C., Zent, R., et al. (2014). Glomerular cell cross-talk influences composition and assembly of extracellular matrix. J. Am. Soc. Nephrol. 25, 953–966.

Chew, C., and Lennon, R. (2018). Basement membrane defects in genetic kidney diseases. Front. Pediatr. 6, 11.

Chiotaki, R., Petrou, P., Giakoumaki, E., Pavlakis, E., Sitaru, C., and Chalepakis, G. (2007). Spatiotemporal distribution of Fras1/Frem proteins during mouse embryonic development. Gene Expr Patterns 7, 381–388.

Combes, A.N., Zappia, L., Er, P.X., Oshlack, A., and Little, M.H. (2019a). Single-cell analysis reveals congruence between kidney organoids and human fetal kidney. Genome Med. 11, 3.

Combes, A.N., Phipson, B., Lawlor, K.T., Dorison, A., Patrick, R., Zappia, L., Harvey, R.P., Oshlack, A., and Little, M.H. (2019b). Single cell analysis of the developing mouse kidney provides deeper insight into marker gene expression and ligand-receptor crosstalk. Development 146.

Czerniecki, S.M., Cruz, N.M., Harder, J.L., Menon, R., Annis, J., Otto, E.A., Gulieva, R.E., Islas, L.V., Kim, Y.K., Tran, L.M., et al. (2018). High-Throughput Screening Enhances Kidney Organoid Differentiation from Human Pluripotent Stem Cells and Enables Automated Multidimensional Phenotyping. Cell Stem Cell 22, 929–940.e4.

Feitosa, N.M., Zhang, J., Carney, T.J., Metzger, M., Korzh, V., Bloch, W., and Hammerschmidt, M. (2012). Hemicentin 2 and Fibulin 1 are required for epidermal-dermal junction formation and fin mesenchymal cell migration during zebrafish development. Dev. Biol. 369, 235–248.

Forbes, T.A., Howden, S.E., Lawlor, K., Phipson, B., Maksimovic, J., Hale, L., Wilson, S., Quinlan, C., Ho, G., Holman, K., et al. (2018). Patient-iPSC-Derived Kidney Organoids Show Functional Validation of a Ciliopathic Renal Phenotype and Reveal Underlying Pathogenetic Mechanisms. Am. J. Hum. Genet. 102, 816–831.

Gatseva, A., Sin, Y.Y., Brezzo, G., and Van Agtmael, T. (2019). Basement membrane collagens and disease mechanisms. Essays Biochem 63, 297–312.

Hale, L.J., Howden, S.E., Phipson, B., Lonsdale, A., Er, P.X., Ghobrial, I., Hosawi, S., Wilson, S., Lawlor, K.T., Khan, S., et al. (2018). 3D organoid-derived human glomeruli for personalised podocyte disease modelling and drug screening. Nat. Commun. 9, 5167.

Hebert, J.D., Myers, S.A., Naba, A., Abbruzzese, G., Lamar, J.M., Carr, S.A., and Hynes, R.O. (2020). Proteomic profiling of the ECM of xenograft breast cancer metastases in different organs reveals distinct metastatic niches. Cancer Res. 80, 1475–1485.

Howard, A.M., LaFever, K.S., Fenix, A.M., Scurrah, C.R., Lau, K.S., Burnette, D.T., Bhave, G., Ferrell, N., and Page-McCaw, A. (2019). DSS-induced damage to basement membranes is repaired by matrix replacement and crosslinking. J. Cell Sci. 132.

Jayadev, R., and Sherwood, D.R. (2017). Basement membranes. Curr. Biol. 27, R207–R211.

Jayadev, R., Chi, Q., Keeley, D.P., Hastie, E.L., Kelley, L.C., and Sherwood, D.R. (2019). α-Integrins dictate distinct modes of type IV collagen recruitment to basement membranes. J. Cell Biol. 218, 3098–3116.

Kashtan, C.E., Kim, Y., Lees, G.E., Thorner, P.S., Virtanen, I., and Miner, J.H. (2001). Abnormal glomerular basement membrane laminins in murine, canine, and human Alport syndrome: aberrant laminin alpha2 deposition is species independent. J. Am. Soc. Nephrol. 12, 252–260.

Keeley, D.P., Hastie, E., Jayadev, R., Kelley, L.C., Chi, Q., Payne, S.G., Jeger, J.L., Hoffman, B.D., and Sherwood, D.R. (2020). Comprehensive Endogenous Tagging of Basement Membrane Components Reveals Dynamic Movement within the Matrix Scaffolding. Dev. Cell 54, 60–74.e7.

Kruegel, J., and Miosge, N. (2010). Basement membrane components are key players in specialized extracellular matrices. Cell Mol. Life Sci. 67, 2879–2895.

Kyprianou, C., Christodoulou, N., Hamilton, R.S., Nahaboo, W., Boomgaard, D.S., Amadei, G., Migeotte, I., and Zernicka-Goetz, M. (2020). Basement membrane remodelling regulates mouse embryogenesis. Nature 582, 253–258.

Lennon, R., Byron, A., Humphries, J.D., Randles, M.J., Carisey, A., Murphy, S., Knight, D., Brenchley, P.E., Zent, R., and Humphries, M.J. (2014). Global analysis reveals the complexity of the human glomerular extracellular matrix. J. Am. Soc. Nephrol. 25, 939–951.

Li, S., Edgar, D., Fässler, R., Wadsworth, W., and Yurchenco, P.D. (2003). The role of laminin in embryonic cell polarization and tissue organization. Dev. Cell 4, 613–624.

Lipp, S.N., Jacobson, K.R., Hains, D.S., Schwarderer, A.L., and Calve, S. (2021). 3D mapping reveals a complex and transient interstitial matrix during murine kidney development. J. Am. Soc. Nephrol.

Maier, J.I., Rogg, M., Helmstädter, M., Sammarco, A., Schilling, O., Sabass, B., Miner, J.H., Dengjel, J., Walz, G., Werner, M., et al. (2021). EPB41L5 controls podocyte extracellular matrix assembly by adhesome-dependent force transmission. Cell Rep. 34, 108883.

Matsubayashi, Y., Louani, A., Dragu, A., Sánchez-Sánchez, B.J., Serna-Morales, E., Yolland, L., Gyoergy, A., Vizcay, G., Fleck, R.A., Heddleston, J.M., et al. (2017). A moving source of matrix components is essential for de novo basement membrane formation. Curr. Biol. 27, 3526–3534.e4.

Matsubayashi, Y., Sánchez-Sánchez, B.J., Marcotti, S., Serna-Morales, E., Dragu, A., Díaz-de-la-Loza, M.-D.-C., Vizcay-Barrena, G., Fleck, R.A., and Stramer, B.M. (2020). Rapid Homeostatic Turnover of Embryonic ECM during Tissue Morphogenesis. Dev. Cell 54, 33–42.e9.

Miner, J.H. (1999). Renal basement membrane components. Kidney Int. 56, 2016–2024.

Miner, J.H., and Sanes, J.R. (1996). Molecular and functional defects in kidneys of mice lacking collagen alpha 3(IV): implications for Alport syndrome. J. Cell Biol. 135, 1403–1413.

Miner, J.H., and Yurchenco, P.D. (2004). Laminin functions in tissue morphogenesis. Annu. Rev. Cell Dev. Biol. 20, 255–284.

Morello, R., Zhou, G., Dreyer, S.D., Harvey, S.J., Ninomiya, Y., Thorner, P.S., Miner, J.H., Cole, W., Winterpacht, A., Zabel, B., et al. (2001). Regulation of glomerular basement membrane collagen expression by LMX1B contributes to renal disease in nail patella syndrome. Nat. Genet. 27, 205–208.

Morrissey, M.A., Keeley, D.P., Hagedorn, E.J., McClatchey, S.T.H., Chi, Q., Hall, D.H., and Sherwood, D.R. (2014). B-LINK: a hemicentin, plakin, and integrin-dependent adhesion system that links tissues by connecting adjacent basement membranes. Dev. Cell 31, 319–331.

Naba, A., Clauser, K.R., Ding, H., Whittaker, C.A., Carr, S.A., and Hynes, R.O. (2016). The extracellular matrix: Tools and insights for the “omics” era. Matrix Biol 49, 10–24.

Naba, A., Pearce, O.M.T., Del Rosario, A., Ma, D., Ding, H., Rajeeve, V., Cutillas, P.R., Balkwill, F.R., and Hynes, R.O. (2017). Characterization of the extracellular matrix of normal and diseased tissues using proteomics. J. Proteome Res. 16, 3083–3091.

Naylor, R.W., Morais, M., and Lennon, R. (2020). Complexities of the glomerular basement membrane. Nat. Rev. Nephrol. 17, 112–127.

Petrou, P., Makrygiannis, A.K., and Chalepakis, G. (2008). The Fras1/Frem family of extracellular matrix proteins: structure, function, and association with Fraser syndrome and the mouse bleb phenotype. Connect Tissue Res 49, 277–282.

Randles, M.J., Woolf, A.S., Huang, J.L., Byron, A., Humphries, J.D., Price, K.L., Kolatsi-Joannou, M., Collinson, S., Denny, T., Knight, D., et al. (2015). Genetic background is a key determinant of glomerular extracellular matrix composition and organization. J. Am. Soc. Nephrol. 26, 3021–3034.

Randles, M.J., Humphries, M.J., and Lennon, R. (2017). Proteomic definitions of basement membrane composition in health and disease. Matrix Biol 57–58, 12–28.

Romero-Guevara, R., Ioannides, A., and Xinaris, C. (2020). Kidney organoids as disease models: strengths, weaknesses and perspectives. Front. Physiol. 11, 563981.

Rooney, K.M., Woolf, A.S., and Kimber, S.J. (2021). Towards Modelling Genetic Kidney Diseases with Human Pluripotent Stem Cells. Nephron 145, 285–296.

Sasaki, T., Fässler, R., and Hohenester, E. (2004). Laminin: the crux of basement membrane assembly. J. Cell Biol. 164, 959–963.

Takasato, M., Er, P.X., Chiu, H.S., Maier, B., Baillie, G.J., Ferguson, C., Parton, R.G., Wolvetang, E.J., Roost, M.S., Chuva de Sousa Lopes, S.M., et al. (2015). Kidney organoids from human iPS cells contain multiple lineages and model human nephrogenesis. Nature 526, 564–568.

Tanigawa, S., Islam, M., Sharmin, S., Naganuma, H., Yoshimura, Y., Haque, F., Era, T., Nakazato, H., Nakanishi, K., Sakuma, T., et al. (2018). Organoids from Nephrotic Disease-Derived iPSCs Identify Impaired NEPHRIN Localization and Slit Diaphragm Formation in Kidney Podocytes. Stem Cell Rep. 11, 727–740.

Tian, P., and Lennon, R. (2019). The myriad possibility of kidney organoids. Curr Opin Nephrol Hypertens 28, 211–218.

Urbano, J.M., Torgler, C.N., Molnar, C., Tepass, U., López-Varea, A., Brown, N.H., de Celis, J.F., and Martín-Bermudo, M.D. (2009). Drosophila laminins act as key regulators of basement membrane assembly and morphogenesis. Development 136, 4165–4176.

Wu, H., Uchimura, K., Donnelly, E.L., Kirita, Y., Morris, S.A., and Humphreys, B.D. (2018). Comparative Analysis and Refinement of Human PSC-Derived Kidney Organoid Differentiation with Single-Cell Transcriptomics. Cell Stem Cell 23, 869–881.e8.

Yamamura, T., Horinouchi, T., Nagano, C., Omori, T., Sakakibara, N., Aoto, Y., Ishiko, S., Nakanishi, K., Shima, Y., Nagase, H., et al. (2020a). Genotype-phenotype correlations influence the response to angiotensin-targeting drugs in Japanese patients with male X-linked Alport syndrome. Kidney Int. 98, 1605–1614.

Yamamura, T., Horinouchi, T., Adachi, T., Terakawa, M., Takaoka, Y., Omachi, K., Takasato, M., Takaishi, K., Shoji, T., Onishi, Y., et al. (2020b). Development of an exon skipping therapy for X-linked Alport syndrome with truncating variants in COL4A5. Nat. Commun. 11, 2777.

Young, M.D., Mitchell, T.J., Vieira Braga, F.A., Tran, M.G.B., Stewart, B.J., Ferdinand, J.R., Collord, G., Botting, R.A., Popescu, D.-M., Loudon, K.W., et al. (2018). Single-cell transcriptomes from human kidneys reveal the cellular identity of renal tumors. Science 361, 594–599.

Zenker, M., Aigner, T., Wendler, O., Tralau, T., Müntefering, H., Fenski, R., Pitz, S., Schumacher, V., Royer-Pokora, B., Wühl, E., et al. (2004). Human laminin beta2 deficiency causes congenital nephrosis with mesangial sclerosis and distinct eye abnormalities. Hum. Mol. Genet. 13, 2625–2632.

