## Supplemental information for "Kidney organoids: A system to study human basement membrane assembly in health and disease"

*1.Wellcome Centre for Cell-Matrix Research, Division of Cell-Matrix Biology and Regenerative Medicine, School of Biological Sciences, Faculty of Biology Medicine and Health, The University of Manchester, Manchester Academic Health Science Centre, Manchester, M13 9PT, UK. 2. Division of Informatics, Imaging & Data Sciences, School of Biological Sciences, Faculty of Biology Medicine and Health, The University of Manchester, Manchester Academic Health Science Centre, Manchester, M13 9PT, UK. 3. Division of Cell Matrix Biology & Regenerative Medicine, School of Biological Sciences, Faculty of Biology Medicine and Health, The University of Manchester, Manchester Academic Health Science Centre, Manchester, M13 9PT, UK. 4. Electron Microscopy Core Facility, RRID:SCR_021147, Faculty of Biology, Medicine and Health, The University of Manchester, Manchester, M13 9PT, UK. 5. Developmental Biology and Cancer Programme, UCL Great Ormond Institute of Child Health, London, WC1N 1EH, UK. 6. Department of Renal Medicine, University College London, London, NW3 2PF, UK. 7. Department of Cell and Developmental Biology, Institute of Biomedical Sciences, University of São Paulo, São Paulo, SP, 05508-000, Brazil. 8. Department of Medicine, Vanderbilt University Medical Center, Nashville, TN, 37232, USA. 10. Department of Paediatric Nephrology, Royal Manchester Children's Hospital, Manchester University Hospitals NHS Foundation Trust, Manchester Academic Health Science Centre, Manchester, M13 9WL, UK.*

**Author list footnotes**:

**Supplemental information**

### Clinical presentation

The index case presented aged 4, with recurrent episodes of macroscopic haematuria and persistent microscopic haematuria. His mother also exhibited microscopic haematuria and he underwent a kidney biopsy. This had normal appearances by light microscopy and immunohistochemical staining did not show glomerular deposition of immuno-reactants. With electron microscopy the glomerular basement membrane (GBM) was thinned in some glomerular capillary loops. In one or two others the GBM was irregularly thickened with lamination. There were also electron-lucent lacunae between the laminations, some of which contained electron-dense regions and the ultrastructural changes, in combination with the highly suggestive presentation and family history, confirmed a diagnosis of Alport Syndrome. The patient subsequently developed hypertension and was treated with antihypertensive medications, including renin-angiotensin system blockade, and exhibited slow decline in his kidney function. He eventually progressed to end stage kidney disease and received a pre-emptive kidney transplant from his healthy father at the age of 19. Subsequent genetic testing identified a likely pathogenic missense variant in *COL4A5* (c.3595G>A; p.Gly1232Asp) and a variant of unknown significance in *COL4A4* (c.3286C>T; p.Pro1096Ser). Further genetic testing in the family confirmed that both the patient's mother and sister carried one or both of these variants. His mother had both the *COL4A5* and *COL4A4* variants and exhibited persistent microscopic haematuria and urine protein:creatinine ratios (uPCR) ranging from 17 to 30 mg/mmol and estimated glomerular filtration rate steady between 74 to 85 ml/min/1.73m$^2$ body surface area over 5 years to most recent follow-up aged 55. The index patient's sister shared only the *COL4A5* variant and also exhibited persistent haematuria with proteinuria (uPCR 264 mg/mmol falling to 45-89 mg/mmol on institution of renin-angiotensin system blockade aged 25). Serum creatinine was normal with eGFR >90 ml/min/1.73m$^2$ body surface area at last follow-up aged 30. The *COL4A4* variant was also detected in the index case's healthy maternal grandmother (in whom urinalysis was normal) but the *COL4A5* variant was absent from both maternal grandparents, suggesting a *de novo* variant in his mother.

### METHODS

#### Antibodies

For immunofluorescence (IF) and whole mount immunofluorescence (WM), we used the following primary antibodies: mouse mAb anti-CD31 [89C2] (3528, Cell Signaling) diluted 1:100 (IF, WM), mouse mAb anti-E cadherin [M168] (ab76055, Abcam) diluted 1:300 (IF, WM), rabbit pAb anti-WT1 [C-19] (sc-192, Santa Cruz Biotechnology) diluted 1: 100 (IF, WM), sheep pAb anti-human nephrin (AF4269, R&D Systems) diluted 1: 400 (WM) or 1:200 (IF), rat mAb anti-human collagen alpha-1(IV) NC1 domain [H11] (7070, Chondrex) diluted 1: 400 (WM) or 1:100 (IF), rat mAb anti-human collagen alpha-3(IV) NC1 domain [H31] (7076, Chondrex) diluted 1: 400 (WM) or 1:100 (IF), rat mAb anti-human collagen alpha-4(IV) NC1 domain [H43] (7073, Chondrex) diluted 1: 400 (WM) or 1:100 (IF), rat mAb anti-human collagen alpha-6(IV) NC1 domain [H63] (7074, Chondrex) diluted 1: 400 (WM) or 1:50 (IF), rabbit pAb anti-laminin (ab11575, Abcam) diluted 1:250 (IF, WM), mouse mAb anti-nidogen [302117] (MA5-23911, Invitrogen) diluted 8.3 μg/ml (IF, WM), rat mAb anti-perlecan [A7L6] (MAB1948P, MilliporeSigma) diluted 1:250 (IF, WM), mouse mAb anti-laminin alpha-5 chain [4C7] (ab17107, Abcam) diluted 1:100 (IF), mouse mAb anti-laminin beta-1 chain [4E10] (MAB1921P, MilliporeSigma) diluted 1:100 (IF), mouse mAb anti-laminin S/laminin beta 2 chain [CL2979] (NBP2-42387, Novus Biologicals) diluted 1:50 (IF), rabbit pAb anti-podocin antibody (P0372, MilliporeSigma) diluted 1:200 (IF), rabbit anti-NPHS2 (ab50339, Abcam) diluted 1:200 (IF); secondary antibodies were: donkey anti-rat IgG conjugated with Alexa Fluor 488 (A21208, Invitrogen Antibodies) or Alexa Fluor 594 (A-21209, Invitrogen Antibodies) diluted 1:400, donkey anti-mouse IgG conjugated with Alexa Fluor 488 (A21202, Invitrogen Antibodies) or Alexa Fluor 594 (A21203, Invitrogen Antibodies) diluted 1:400, donkey anti-rabbit IgG conjugated with Alexa Fluor 488 (A21206, Invitrogen Antibodies) or Alexa Fluor 647 (A31573, Invitrogen Antibodies) diluted 1:400, goat anti-rabbit IgG conjugated with Alexa Fluor 594 (A11012, Invitrogen Antibodies) diluted 1:400, donkey anti-sheep IgG conjugated with Alexa Fluor 594 (A11016, Invitrogen Antibodies) or Alexa Fluor 680 (A21102, Invitrogen Antibodies) diluted 1:400. For immunoblotting, we used a mouse mAb anti-laminin S/laminin beta 2 chain [CL2979] (NBP2-42387, Novus Biologicals) diluted 1:1000, and a secondary goat anti-mouse IgG DyLight 800 4X PEG conjugated (5151S, Cell Signaling Technology) diluted 1:1000.

**Human fetal kidney**

Human fetal kidney sections were provided by the Joint MRC/Wellcome Trust Human Developmental Biology Resource (HDBR) (http://hdbr.org). The HDBR obtains written consent from the donors and has ethics approval (REC reference: 08/H0712/34+5) to collect, store and distribute human material sampled between 4- and 21-weeks post conception. All experimental protocols were approved by the Institute's Ethical Committee (reference 010/H0713/6) and were performed in accordance with institutional ethical and regulatory guidelines.

**Induced pluripotent stem cells**

Human induced pluripotent stem cells (iPSCs) derived from peripheral blood mononuclear cells (PBMCs) of a healthy individual were generated as previously described (Wood et al.,

2020). Briefly, PBMCs were isolated from whole blood using Ficoll-Paque (GE17-1440, GE Healthcare). Isolated PBMCs were then grown in StemSpan Erythroid Expansion Medium (02692, Stem Cell Technologies) for 8 days before being transduced using CytoTune-iPS 2.0 Sendai virus (A16517, Invitrogen). Transduced cells where then grown on vitronectin (A14700, Gibco) coated plates in ReproTeSR medium (05926, Stem Cell Technologies). When large enough, colonies were isolated by manual picking and grown on vitronectin coated plates in TeSR-E8 medium (05991, STEMCELL). iPSCs derived from patients with Alport syndrome were generated at the Wellcome Sanger Institute, in collaboration with the Human Induced Pluripotent Stem Cell Initiative (HipSci, www.hipsci.org; (Kilpinen et al., 2017). Following patient consent, dermal fibroblasts were obtained from skin biopsies and programmed to iPSC. For this study we studied three members of the same family. A male index case and his mother both with a likely pathogenic *COL4A5* variant (c.3595G>A; p.Gly1232Asp) and a *COL4A4* variant of uncertain significance (c.3286C>T; p.Pro1096Ser) and the sister of the index case carrying only the *COL4A5* variant.

**Kidney organoid differentiation**

We used iPSCs between passage 20 and 30 and differentiated these cells to kidney organoids as previously described (Takasato et al., 2015). Briefly iPSCs were maintained at 37°C with TeSR™-E8 medium with 25x Supplement (05991, 05992, STEMCELL) in 6-well plates (3516, Corning) coated with vitronectin (A14700, Gibco). Prior to differentiation (day 0) cells were dissociated with TrypLE (12563029, Thermo Fisher Scientific), counted with a hemocytometer and seeded in vitronectin-coated 24-well plates (3524, Corning) at a density of 35,000 cells/cm$^2$ in TeSR™-E8™ medium with Revitacell 10 µl/ml (A2644501, Gibco). Intermediate mesoderm induction was performed by changing medium after 24 hours to STEMdiff™ APEL™2 (05270, STEMCELL™) with 3% protein free hybridoma medium (12040077, Gibco), and 8 µM CHIR99021 (4423/10, Tocris Bioscience) for 4 days. On day 5, CHIR99021 was replaced by 200 ng/ml FGF-9 (100-23, Peprotech) and 1 µg/ml heparin (H3393, Sigma Aldrich) for a further 7 days. On Day 7, cells were dissociated with TrypLE, counted and pelletized into organoids (250,000 cells) by centrifuging 400x g/min four times. Organoids were carefully placed on 0.4 µm Millicell Cell Culture Insert in 6-well plates (PICM0RG50, Millipore), and were cultured for 1 hour in the presence of 5 µM CHIR99021 in APEL™2 medium, then subsequently cultured in APEL™2 medium supplemented with 200 ng/mL FGF9 and 1 µg/mL Heparin until Day 11. From day 12, organoids were grown in STEMdiff™ APEL™2 without growth factors, with medium change every 2 days.

**Mice**

All mouse handling and experimental procedures were approved by the Animal Ethics Committee of the Institute of Biomedical Sciences (University of São Paulo, Brazil; reference 019/2015). 2-month-old *Swiss* female mice were housed in the Experimental Animal Facility (ICB, USP) kept in groups of 3-4 subjects per cage (41x34x16 cm) at 12-hour light/dark cycle at 25 °C, with free access to water and chow. Mating occurred overnight, and females were checked for vaginal plugs on the next morning to determine if mating had occurred and gestation was timed accordingly (embryonic day 1). Pregnant dams were separate and kept in individual cages (30x2013 cm) under the same conditions above mentioned. Fetuses were collected on the embryonic day 19 (E19), following C-section surgery in the pregnant mice under anaesthesia with 25 mg/kg avertin (T48402, Sigma-Aldrich). Fetal kidneys were dissected and processed for histological analysis, or snap-frozen in liquid nitrogen for proteomic analysis.

**Whole mount immunofluorescence**

Whole organoids were fixed with 2% (wt/vol) paraformaldehyde at 4°C for 20 minutes, washed with Phosphate Buffered Saline (PBS; D8537, Sigma-Aldrich) and blocked with 1x casein blocking buffer (B6429, SigmaAldrich) for 2 hours at room temperature. Samples were incubated overnight with primary antibodies in blocking buffer at 4°C. After thoroughly washing with 0.3% (vol/vol) Triton X-100 in PBS (PBS-Triton; X100, SigmaAldrich), the samples were incubated overnight with Alexa Fluor-conjugated secondary antibodies at 4°C. Nuclei were stained with Hoechst 33342 solution (B2261, SigmaAldrich). Imaging was performed in glass-bottomed dishes (P35G-1.5-10-C, MatTek) mounted with ProLong Gold Antifade mountant (P36934, Invitrogen). Images were acquired with a Leica TCS SP8 AOBS inverted confocal using a 63x HCX PL APO (Oil) objective, using hybrid detectors with the following detection mirror settings: FITC 494-530 nm; Texas red 602-665 nm; Cy5 640-690 nm. When it was not possible to eliminate fluorescence crosstalk, the images were collected sequentially. When acquiring 3D optical stacks, the confocal software was used to determine the optimal number of Z sections. Only the maximum intensity projections of these 3D stacks are shown in the results. 3D image stacks were analyzed with ImageJ v 1.53g software (Rasband, W.S., ImageJ, U. S. National Institutes of Health, Bethesda, Maryland, USA; available at https://imagej.nih.gov/ij/, 1997-2018).

**Histology and immunofluorescence**

For light microscopy, formaldehyde-fixed paraffin-embedded samples were cut and stained with hematoxylin and eosin (H&E) for morphological analysis. Images were acquired on a 3D-Histech Pannoramic-250 microscope slide-scanner using a x20 objective (Zeiss) using the Case Viewer software (3D-Histech). For immunofluorescence microscopy, 8-μm-thick sections were air dried and fixed with cold acetone, or alternatively, 5-μm-thick paraffin sections underwent antigen retrieval with 10 mM sodium citrate buffer (pH 2.0 or 6.0) in a microwave for 15 min at 30% power, and further treated with 0.1M glycine/6M urea solution for 30 minutes at room temperature. Following nonspecific binding site blocking with 10% normal donkey serum in 1% BSA/PBS, sections were incubated overnight with primary antibody solutions at 4°C. Species-specific secondary antibodies conjugated with either Alexa Fluor or HRP were used, and nuclei were stained with Hoechst 33342 and mounted with ProLong Gold Antifade mountant. The slides were analyzed with a Zeiss Axioimager.D2 upright microscope equipped with a Coolsnap HQ2 camera (Photometrics) Images were acquired with the Micromanager Software v1.4.23 processed using ImageJ.

**Electron microscopy**

Whole mount primary antibody labelling was performed as described above. After overnight incubation at 4 °C with rabbit pAb anti-laminin (ab11575, Abcam) diluted 1:250, organoids were washed with PBS-Triton and then incubated overnight at 4 °C with a goat anti-rabbit IgG labelled with 10nm gold (ab39601, Abcam) diluted 1:400. After washing with PBS-Triton, the organoids were Stored with 4% paraformaldehyde and 2.5% (wt/vol) glutaraldehyde (Agar Scientific, UK) in 0.1 M HEPES (H0887, SigmaAldrich) pH 7.2, and postfixed with 1% osmium tetroxide (R1024, Agar Scientific) and 1.5% potassium ferrocyanide (214022, The British Drug House, Laboratory Chemicals Division) in 0.1M cacodylate buffer (ph7.2) (R1102, Agar Scientific) for 1 hour then in 1% uranyl acetate (R1100A, Agar Scientific) in water for overnight. After that, the samples were dehydrated in ethanol series infiltrated with low viscosity medium grade resin (T262, TAAB Laboratories Equipment Ltd) and polymerized for 24 hours at 60 °C. For routine TEM sections were cut with Reichert Ultracut ultramicrotome and observed with FEI Tecnai 12 Biotwin microscope at 80kV accelerating voltage. Images were taken with Gatan Orius SC1000 CCD camera.

**SDS-PAGE and immunoblotting**

Whole organoids were harvested and homogenized in ice cold Pierce™ IP Lysis Buffer Proteins (87787, ThermoFisher) with protease inhibitor (04693159001, Roche). The lysates were centrifuged at 4300xg at 4°C for 20 minutes. The supernatant was collected and denatured at 95°C for 5 minutes. Proteins were resolved by SDS-PAGE in a NuPAGE 4-12% Bis-Tris gel (NP0322, Invitrogen) using 1x MOPS SDS running buffer (NP0001, Thermo-Fisher Scientific), and wet-transferred to a nitrocellulose membrane (Z612391, Whatman). Gel loading was assessed by Ponceau S staining (P7170, SigmaAldrich). The membrane blocked with Odyssey blocking buffer in TBS (927-40000, LI-COR) and probed with mouse anti-laminin beta 2 antibody (NBP2-42387, Novus Biologicals) diluted 1:1000, washed incubated with fluorescent-conjugated goat anti-mouse antibody (5151S, Cell Signaling Technology) diluted 1:1000. Bands were visualized using the Odyssey CLx imaging system (LI-COR Biosciences), and background-corrected band optical densitometry was determined using ImageJ.

**Sample enrichment for proteomics analysis**

Kidney organoids from day 14, 18 and 25 of differentiation (*n*=3 per time point) and E19 mouse fetal kidneys were enriched for matrix proteins as previously described (Lennon et al., 2014). Briefly, samples were manually homogenized and incubated in Tris-buffer (10 mM Tris pH 8.0, 150 mM NaCl, 25 mM EDTA, 1% Triton X-100 and 1x with EDTA-free protease inhibitor cocktail (04-693-159-001, Roche) for 1h to extract soluble proteins. After centrifuging (at 14,000x *g* for 10 min) the supernatant (fraction 1) was collected and the remaining pellet was resuspended in alkaline detergent buffer (20 mM $Na_4OH$, 0.5% in PBS-Triton) and incubated for 1h to solubilize and disrupt cell-matrix interactions. After centrifuging, the supernatant (fraction 2) was collected and the pellet was resuspended in benzonase buffer (0.4 µg benzonase; E1014-25KU, Sigma-Aldrich) in PBS supplemented with $CaCl_2$ and $MgCl_2$ (D8537, Sigma-Aldrich) and incubated for 30 min at RT to remove DNA/RNA contaminants. Benzonase inactivation was then carried out at 65 °C for 20 min, and after centrifuging, the supernatant (fraction 3) was collected. The remaining pellet was resuspended in a 5x reducing sample buffer (100 mM Tris pH 6.8, 25% glycerol, 10% SDS, 10% β-mercaptoethanol, 0.1% bromophenol blue) to yield the ECM fraction. Fractions 1 to 3 were diluted (1:4) with a 4x sample buffer, and fractions 1 and 2 were combined (1:1) into a cellular fraction. All steps described were carried out on ice to reduce proteolysis and the fractions obtained were stored at 4°C.

**Laser microdissection microscopy**

Mouse kidney samples (E19, *n*=4) were embedded in OCT for cryosectioning. 10-µm-thick cryosections were acquired and placed onto MMI membrane slides (50102, Molecular Machines and Industries), then fixed with 70% ethanol and hydrated to be stained with haematoxylin and eosin. After rinsing in deionized water, the sections were dehydrated and air-dried just before laser cutting. Maturing mouse glomeruli were identified using a 40x/0.5 FL N objective and laser dissected around the Bowman's capsule using an Olympus IX83 Inverted fluorescence snapshot microscopy equipped with MMI CellCut Microdissection system and the MMI CellTools software v.5.0 (Molecular Machines and Industries). Laser settings were speed = 25 µm/s, focus = 16.45 µm, power = 72.5%. A total of 150 glomerular sections per sample were collected onto sticky 0.5 ml microtube caps (Molecular Machines and Industries) and stored at -80°C until further processing for MS-based proteomics.

**Trypsin digestion**

Fractionated samples for mass spectrometry analysis underwent trypsin digestion either in-gel (mouse samples) or in-solution (organoids and laser-captured mouse samples). For the in-gel digestion, samples were run into SDS-PAGE for 2-3 min to concentrate proteins in the gel top, and then further stained with Expedeon InstantBlue (Z2, Fisher Scientific). After destaining with deionized water, gel-top protein bands were excised and cut into small cubes using sterile razor blades, and transferred to a V-bottomed perforated 96-wells plate (Proxeon) to be additionally destained with 50% acetonitrile (ACN) in digestion buffer (25 mM $NaHCO_3$) for 30 min at RT. After centrifuging (1500 rpm for 2 min), gel pieces were shrunk with 50% ACN for 5 min at RT and completely dried by vacuum centrifugation for 20 min at -120 °C. Next gel pieces were rehydrated and reduced with 10 mM dithiothreitol (DTT; D5545, Sigma-Aldrich) in digestion buffer for 1h at 56 °C, then and alkylated with 55 mM iodoacetamide (IA; I149, Sigma-Aldrich) in digestion buffer for 45 min at RT in the dark. After centrifuging, gel pieces were washed and shrunk with alternating incubations with digestion buffer and 100% ACN, and completely dried by vacuum centrifugation. Samples were incubated overnight in digestion buffer containing 1.25 ng/l sequencing grade modified trypsin (V5111, Promega) at 37 °C to allow protein digestion. Peptides were then collected by centrifugation and remaining peptides in the gel pieces were extracted with ACN/formic acid solution and collected by centrifugation. Pooled extracts were dried by vacuum centrifugation, concentrated to 20 µl of 50% ACN in 0.1% formic acid. For the in-solution digestion, samples were sonicated in S-Trap lysis buffer (5% SDS in 50 mM TEAB pH 7.5) using a Covaris LE220+ Focused Ultrasonicator (Covaris). Extracted proteins were reduced with 5 mM DTT at 60 °C for 10 min under agitation (at 800 rpm), then alkylated with 15 mM IA for 10 min at RT in the dark and had 5 mM DTT added to a 10 mM final concentration to quench residual alkylation reaction. Samples were centrifuged for 10 min at 1400 x *g*, then acidified with 1.2% formic acid and transferred to S-Trap Micro Spin columns (Protifi) equipped with collecting 2.0 ml plastic microtubes to retain the flowthrough. Detergents and salt contaminants were removed by centrifugation (4000 x g for 2 min) and proteins trapped in the S-trap resin were digested for 1h at 47°C with 0.12 g/l trypsin prepared in 50 mM TEAB buffer. Peptides were washed with 50 mM TEAB, centrifuged, washed with 0.1% formic acid, centrifuged again, and finally eluted from the S-trap columns with 30% ACN in 0.1% formic acid solution.

**Offline peptide desalting**

Peptide samples were incubated for 5 min at RT with 5.0 mg Oligo™ R3 reverse-phase beads (1133903, Applied Biosystems) prepared in 50% ACN, in a 96-well plate equipped with 0.2 m polyvinylidene fluoride (PVDF) membrane filter (3504, Corning). After centrifuging, the bead-bound peptides were washed twice with 0.1% formic acid, centrifuged again, and eluted from the filter using 30% ACN in 0.1% formic acid. Retrieved peptides were dried by vacuum centrifugation, then resuspended in 5% ACN in 0.1% formic acid and sent to the Bio-MS Core Research Facility (Faculty of Biology, Medicine and Health, University of Manchester) for mass spectrometry analysis.

**Mass spectrometry data acquisition and analysis**

Samples were analyzed by liquid chromatography-tandem mass spectrometry using an UltiMate® 3000 Rapid Separation LC (RSLC, Dionex Corporation, Sunnyvale, CA) coupled to a Q Exactive™ Hybrid Quadrupole-Orbitrap™ (Thermo Fisher Scientific, Waltham, MA) mass spectrometer. Peptides were separated using a gradient from 93% A (0.1% formic acid in water) and 7% B (0.1% formic acid in ACN) to 18% B over 57 min followed by a second gradient to 27% B over 14 min both at 300 nl/min, using a CSH C18 analytical column (Waters). Peptides were selected for fragmentation automatically by data dependent acquisition. Raw spectra data were acquired and analyzed using Proteome Discoverer software v.2.3.0.523 (Thermo Fisher Scientific). MS data was searched against SwissProt and TrEMBL databases (v. 2018_01; OS = *Mus musculus* for mouse samples; OS=*Homo sapiens* for kidney organoids) using SEQUEST HT and Mascot search tools. For the search were considered: tryptic peptides with up to 1 missed cleavage, mass tolerance of 10 ppm for precursor ions and 0.02 Da for fragment ions, carbamidomethylation of cysteine as fixed modification, oxidation of methionine, proline and lysine, and N-terminal acetylation as dynamic modifications. Peptide-spectrum match, peptide and protein false discovery rates were set to 1%, and protein validation was performed using Target/Decoy strategy. Protein abundances were determined by label-free quantification based on precursor ion intensity. Relative changes in protein abundance were determined by calculating abundance ratios accordingly. Finally, results were filtered for significant FDR master proteins identified with $\geq$ 1 unique peptide and detected in 66% of the samples. *The mass spectrometry proteomics data have been deposited to the ProteomeXchange Consortium via the PRIDE partner repository* (Perez-Riverol et al., 2019) *with the dataset identifiers: PXD025838, PXD025874, PXD025911 and PXD026002.*

**Enrichment and interactome analyses**

Gene ontology (GO) enrichment analysis was performed using the DAVID bioinformatics resource v.6.8 ((Huang et al., 2009); available at https://david.ncifcrf.gov), and term enrichment was determined through Fisher's exact test with Benjamini-Hochberg correction, with a term selected as enriched when FDR < 0.1. Pathway enrichment was performed for proteins differentially expressed using the Reactome database ((Jassal et al., 2020); available at https://reactome.org/). To generate interactome figures, a list of proteins was uploaded to STRING v.11.0 (Szklarczyk et al., 2015) to obtain a collection of high confident protein-protien interactions (combined score ≥ 70%), which was further uploaded into Cytoscape v.3.8.1 (Shannon et al., 2003) to customize the interactomes.

**Single cell RNA-sequencing analysis**

We selected three published single cell-RNA sequencing datasets generated from kidney organoids (GSE114802), fetal and adult human kidneys (EGAS00001002325, EGAS00001002553), and fetal mouse kidney (GSE108291) to identify the cellular origins of BM genes. We first removed the low-quality cells from the dataset to ensure that the technical noise do not affect the downstream analysis. We also remove the lowly expressed genes as they do not give much information and are unreliable for downstream statistical analysis (Bourgon et al., 2010). In order to account for the various sequencing depth of each cell, we normalized the raw counts using the deconvolution-based method (Lun et al., 2016a). We then identified the genes that have high variance in their biological component and use these genes for all downstream analysis. We then applied PCA and took the first 14 components of PCA as input to tSNE and used the first 2 components of tSNE to visualize our cells. The cells were then grouped into their putative clusters using the dynamic tree cut method. We used the *findMarkers* function from *Scran* package to identify the marker genes for each of the clusters (Lun et al., 2016b). *findMarkers* uses t-test for the statistical test and reports p-value of the high rank genes that are DE between the group and all other group. These marker genes were then used to manually annotate the cell types of a cluster (see **Table S3** for clustering annotation details). We also applied SingleR to define the cell types based on matched with annotated bulk datasets (Aran et al., 2019). We used the *plotDots* function from scater package to produce the dotplots (McCarthy et al., 2017).

**Statistical analysis**

Statistical analysis was carried out within Proteome Discoverer using an in-built Two-way ANOVA test with post-hoc Benjamini-Hochberg correction. Principal component analysis (PCA) and unsupervised hierarchical clustering based on a Euclidean distance-based complete-linkage matrix were performed using Rstudio (v. 1.2.5042, http://rstudio.com) with the ggplot2 package (v.3.3.2, https://ggplot2.tidyverse.org) that was also used to generate PCA plots and heat maps. For the integrated proteomic analysis, previously published young human glomerular and kidney tubulointerstitial data (PRIDE accession PXD022219) was reprocessed with Proteome Discoverer to allow direct comparisons with newly acquired data. Then, kidney organoid, mouse and human proteomics datasets were compared using Spearman Rank correlation. Dataset comparisons, for both cellular and ECM cellular fractions, were performed separately for the matrisome proteins only and basement membrane proteins only. The ComplexHeatmap package (v2.2.0, (Gu et al., 2016); http://bioconductor.org/packages/release/bioc/html/ComplexHeatmap.html) was used to generate correlation plots.

**Extended data:**

This project contains the following underlying data hosted at:
https://figshare.com/s/f23b9c38103da0009c69

**Figure 1** Original IF Images: **B** Whole-mount immunofluorescence for kidney cell types; **F** Representative whole mount immunofluorescence images of wild-type and Alport kidney organoids; **G** Immunofluorescence for LAMB2.

**Figure 1** Original light microscope Images: **C** Representative photomicrographs of day 18 kidney organoids (left) and human and mouse fetal kidneys (right).

**Figure 1** Original TEM Images: **D** Transmission electron micrographs of tubular BM in day 25 kidney organoid and E19 mouse fetal kidney.

**Figure 1** Original western blotting image: **H** Immunoblotting for LAMB2 using total lysates from wild type and Alport organoids.

**Figure 2** Original IF Images: **A** Confocal immunofluorescence microscopy of wild-type kidney organoids; **B** perlecan and nidogen on days 11, 18 and 25 of differentiation.

**Figure 4** Original IF Images: **A** Immunofluorescence for key type IV collagen and laminin isoforms showing their emergence and distribution in kidney organoid BM; **D** Immunofluorescence for specific collagen IV isoforms in maturing glomeruli in E19 mouse kidney and in glomerular structures (indicated by dashed lines) in day 25 organoids.

**Figure S1. Morphological characteristics of wild-type kidney organoids, fetal human kidney, and Alport kidney organoids.** (A) Bright field images (left) and H&E staining (right) of kidney organoids on day 11, 14, 18 and 25 of differentiation. (B) H&E staining of human fetal kidney at 8 wpc, 12 wpc, 17 wpc, and 21 wpc highlights normal human kidney development. (C) Whole mount immunofluorescence of wild-type and Alport kidney organoids shows comparable deposition of COL4A4 in BM-like structures (arrows) within NPHS1$^+$/CD31$^+$ glomerular cell cluster (g).

**Figure S2. Time course proteomic analysis of kidney organoid differentiation. Related to Figure 2.** (A) Gene ontology (GO) term enrichment analysis for cellular component annotations associated with proteins detected in the cellular and ECM fractions of kidney organoids by mass spectrometry (MS). GO terms were considered enriched when False discovery rate (FDR) < 0.10. (B) Protein interaction network depicts matrisome proteins identified in kidney organoids by MS over the differentiation time course (nodes represent proteins and connecting lines indicate a reported protein-protein interaction). (C) Principal component analysis (PCA) for the matrix proteins identified by MS in the cellular (left plot) and ECM (right plot) fractions of kidney organoids. (D) Top 15 most abundant BM proteins found in kidney organoids by MS. Proteins were ranked according to their normalized abundance levels (LFQ-intensities). Pooled data are shown as median, error bars indicate the 95% confidence interval for the median. (E) Volcano plots show the $\log_2$-fold change (x-axis) versus $-\log_{10}$-*p*-value (y-axis) for proteins differentially expressed in the cellular fraction of kidney organoids from day 14 to 18, and 18 to 25. Key BM proteins significantly up-regulated (FC $\geq$ 1.5, *p*-value < 0.05, Two-way ANOVA test, *n*=3) are indicated (F) Pathway enrichment analysis for proteins (matrix and non-matrix) differentially expressed during kidney organoid differentiation: bar charts depict log-transformed FDR for the top-most enriched pathways (FDR < 0.10). Pathway terms shown were simplified.

**Figure S3. Single cell-RNA sequencing data analysis of human kidney organoids.** (A) Dotplot depicting the cell-specific level of expression of 160 BM gene collection in 25 days kidney organoids. (B) tSNE plot indicates gene expression of *LIMX1b* and *EPB41L5* in 25 days organoids. Arrows indicate podocyte clusters. Data shown in (A) and (B) were obtained from a re-analysis of a publicly available scRNA-seq dataset (GSE114802; (Combes et al., 2019)). (C) Bar charts show the time-dependent changes in protein abundance of EPB41L5 in kidney organoids. Pooled data are shown as median, error bars indicate the 95% confidence interval for the median. Two-way ANOVA test: **p*-value <0.05.

**Figure S4. Proteomic analysis of E19 mouse fetal kidney and correlational comparison with kidney organoid proteomics.** (A) Gene ontology (GO) term enrichment analysis for cellular component annotations associated with proteins detected by MS in the cellular and ECM. (B) GO biological process annotations enriched for top 100 most abundant proteins detected by MS in the cellular and ECM. (C) Top 20 most abundant BM proteins found in the E19 mouse kidney by MS. Proteins were ranked according to their normalized abundance levels (LFQ-intensities). Pooled data are presented as median, error bars indicate the 95% confidence interval for the median. (D) Comparison of other structural matrix and ECM-associated proteins identified in the E19 mouse kidney and kidney organoids over differentiation. (E) Spearman rank correlation plots depicting the r coefficient values for matrix and BM protein abundance (ECM fraction) comparisons between the E19 mouse kidney and kidney organoids.

**Figure S5. Single-cell RNA sequencing analysis of human fetal kidney**. Re-analysis of 8/9-wpc human kidney scRNA-seq datasets (EGAS00001002325, EGAS00001002553; Young et al., 2018) confirms cellular specificity for collagen IV and laminin isoform gene expression. tSNE plots represent the cell type clusters identified, and colour intensity indicate the cell-specific level of expression for the selected BM genes.

**Figure S6 Integrated correlational analysis of organoid and *in vivo* kidney datasets**. Spearman rank correlation plots depicting the r coefficient values for matrix and BM protein abundance (ECM fraction) between E19 mouse kidney, kidney organoids and adult human kidney proteomic datasets.

**Table S1** Human fetal kidney and hiPSC general information

**Table S2** Human kidney organoid proteome and matrix proteins

**Table S3** scRNA-seq kidney datasets- cell clustering and expression data

**Table S4** E19 mouse maturing glomerulus proteome and matrix proteins

**Table S5** E19 mouse kidney proteome and matrix proteins

**Table S6** Human adult kidney glomerular and tubulointerstitial proteome and matrix proteins

**References**

Aran, D., Looney, A.P., Liu, L., Wu, E., Fong, V., Hsu, A., Chak, S., Naikawadi, R.P., Wolters, P.J., Abate, A.R., et al. (2019). Reference-based analysis of lung single-cell sequencing reveals a transitional profibrotic macrophage. Nat. Immunol. *20*, 163–172.

Bourgon, R., Gentleman, R., and Huber, W. (2010). Independent filtering increases detection power for high-throughput experiments. Proc. Natl. Acad. Sci. USA *107*, 9546–9551.

Combes, A.N., Zappia, L., Er, P.X., Oshlack, A., and Little, M.H. (2019). Single-cell analysis reveals congruence between kidney organoids and human fetal kidney. Genome Med. *11*, 3.

Gu, Z., Eils, R., and Schlesner, M. (2016). Complex heatmaps reveal patterns and correlations in multidimensional genomic data. Bioinformatics *32*, 2847–2849.

Huang, D.W., Sherman, B.T., and Lempicki, R.A. (2009). Systematic and integrative analysis of large gene lists using DAVID bioinformatics resources. Nat. Protoc. *4*, 44–57.

Jassal, B., Matthews, L., Viteri, G., Gong, C., Lorente, P., Fabregat, A., Sidiropoulos, K., Cook, J., Gillespie, M., Haw, R., et al. (2020). The Reactome Pathway Knowledgebase. Nucleic Acids Res. *48*, D498–D503.

Kilpinen, H., Goncalves, A., Leha, A., Afzal, V., Alasoo, K., Ashford, S., Bala, S., Bensaddek, D., Casale, F.P., Culley, O.J., et al. (2017). Common genetic variation drives molecular heterogeneity in human iPSCs. Nature *546*, 370–375.

Lennon, R., Byron, A., Humphries, J.D., Randles, M.J., Carisey, A., Murphy, S., Knight, D., Brenchley, P.E., Zent, R., and Humphries, M.J. (2014). Global analysis reveals the complexity of the human glomerular extracellular matrix. J. Am. Soc. Nephrol. *25*, 939–951.

Lun, A.T.L., Bach, K., and Marioni, J.C. (2016a). Pooling across cells to normalize single-cell RNA sequencing data with many zero counts. Genome Biol. *17*, 75.

Lun, A.T.L., McCarthy, D.J., and Marioni, J.C. (2016b). A step-by-step workflow for low-level analysis of single-cell RNA-seq data with Bioconductor. [version 2; peer review: 3 approved, 2 approved with reservations]. F1000Res. *5*, 2122.

McCarthy, D.J., Campbell, K.R., Lun, A.T.L., and Wills, Q.F. (2017). Scater: pre-processing, quality control, normalization and visualization of single-cell RNA-seq data in R. Bioinformatics *33*, 1179–1186.

Perez-Riverol, Y., Csordas, A., Bai, J., Bernal-Llinares, M., Hewapathirana, S., Kundu, D.J., Inuganti, A., Griss, J., Mayer, G., Eisenacher, M., et al. (2019). The PRIDE database and related tools and resources in 2019: improving support for quantification data. Nucleic Acids Res. *47*, D442–D450.

Shannon, P., Markiel, A., Ozier, O., Baliga, N.S., Wang, J.T., Ramage, D., Amin, N., Schwikowski, B., and Ideker, T. (2003). Cytoscape: a software environment for integrated models of biomolecular interaction networks. Genome Res. *13*, 2498–2504.

Szklarczyk, D., Franceschini, A., Wyder, S., Forslund, K., Heller, D., Huerta-Cepas, J., Simonovic, M., Roth, A., Santos, A., Tsafou, K.P., et al. (2015). STRING v10: protein-protein interaction networks, integrated over the tree of life. Nucleic Acids Res. *43*, D447-52.

Takasato, M., Er, P.X., Chiu, H.S., Maier, B., Baillie, G.J., Ferguson, C., Parton, R.G., Wolvetang, E.J., Roost, M.S., Chuva de Sousa Lopes, S.M., et al. (2015). Kidney organoids from human iPS cells contain multiple lineages and model human nephrogenesis. Nature *526*, 564–568.

Wood, K.A., Rowlands, C.F., Thomas, H.B., Woods, S., O'Flaherty, J., Douzgou, S., Kimber, S.J., Newman, W.G., and O'Keefe, R.T. (2020). Modelling the developmental spliceosomal craniofacial disorder Burn-McKeown syndrome using induced pluripotent stem cells. PLoS One *15*, e0233582.

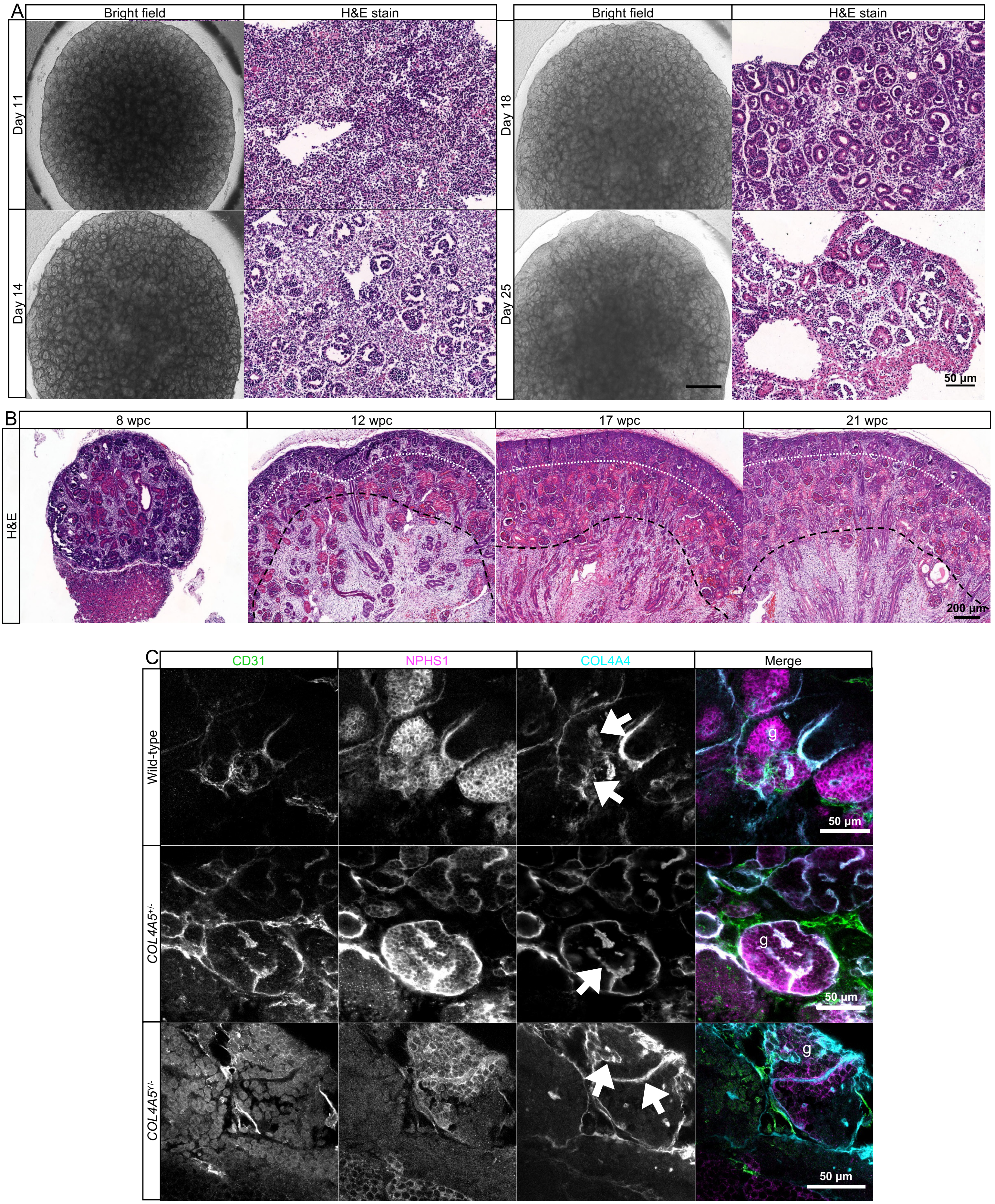
A
Bright field
H&E stain
Bright field
H&E stain
Day 11
Day 14
Day 18
Day 25
50 µm
B
8 wpc
12 wpc
17 wpc
21 wpc
H&E
200 µm
C
CD31
NPHS1
COL4A4
Merge
Wild-type
COL4A5+/-
COL4A5Y/-
g
50 µm

Figure S1

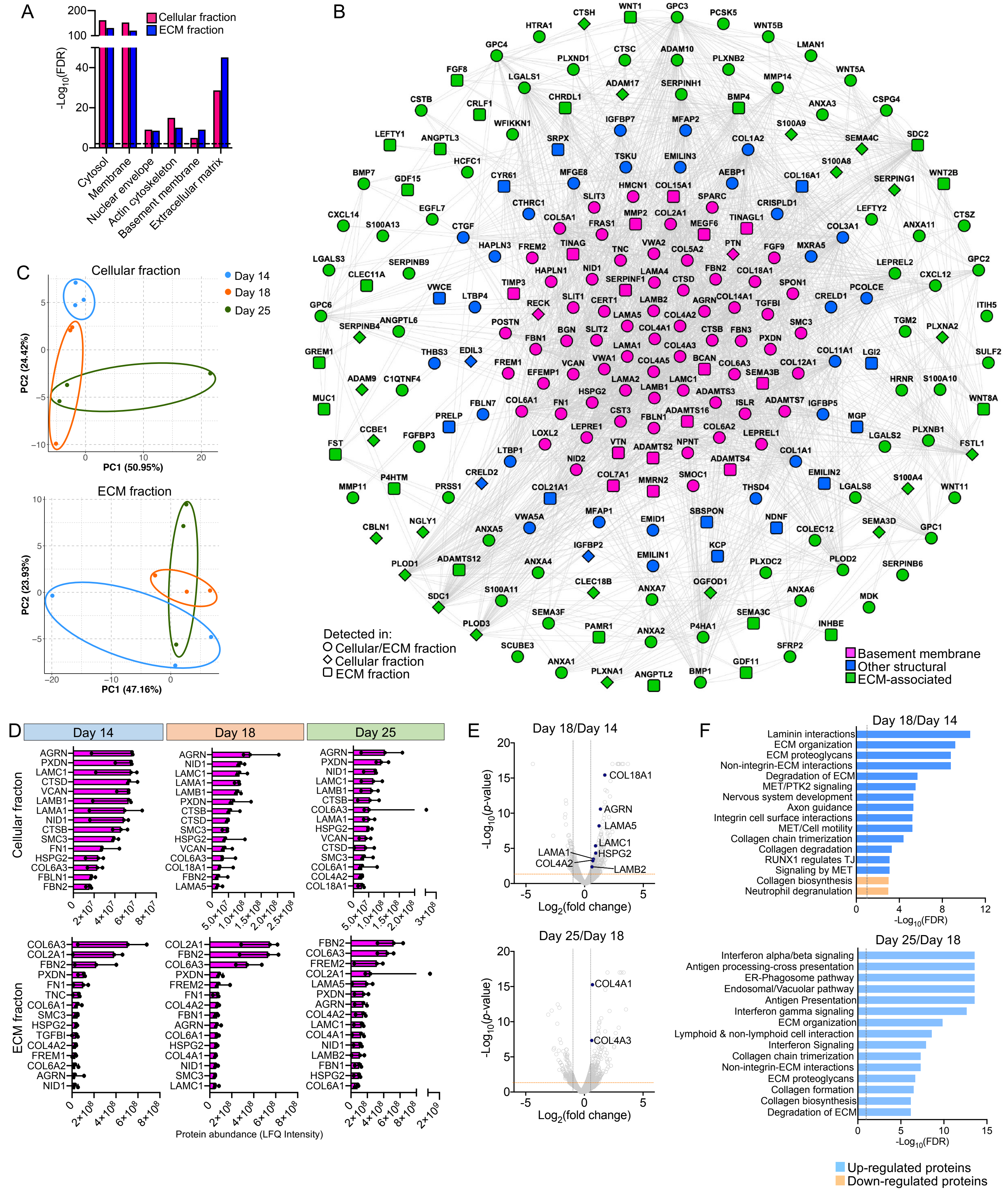

A
Cellular fraction
ECM fraction
-Log10(FDR)
Cytosol
Membrane
Nuclear envelope
Actin cytoskeleton
Basement membrane
Extracellular matrix
B
Detected in:
Cellular/ECM fraction
Cellular fraction
ECM fraction
Basement membrane
Other structural
ECM-associated
C
Cellular fraction
Day 14
Day 18
Day 25
PC1 (50.95%)
PC2 (24.42%)
ECM fraction
PC1 (47.16%)
PC2 (23.93%)
D
Day 14
Day 18
Day 25
Cellular fracton
ECM fracton
Protein abundance (LFQ Intensity)
E
Day 18/Day 14
Day 25/Day 18
-Log10(p-value)
Log2(fold change)
COL18A1
AGRN
LAMA5
LAMC1
HSPG2
LAMA1
COL4A2
LAMB2
COL4A1
COL4A3
F
Day 18/Day 14
Laminin interactions
ECM organization
ECM proteoglycans
Non-integrin-ECM interactions
Degradation of ECM
MET/PTK2 signaling
Nervous system development
Axon guidance
Integrin cell surface interactions
MET/Cell motility
Collagen chain trimerization
Collagen degradation
RUNX1 regulates TJ
Signaling by MET
Collagen biosynthesis
Neutrophil degranulation
-Log10(FDR)
Day 25/Day 18
Interferon alpha/beta signaling
Antigen processing-cross presentation
ER-Phagosome pathway
Endosomal/Vacuolar pathway
Antigen Presentation
Interferon gamma signaling
ECM organization
Lymphoid & non-lymphoid cell interaction
Interferon Signaling
Collagen chain trimerization
Non-integrin-ECM interactions
ECM proteoglycans
Collagen formation
Collagen biosynthesis
Degradation of ECM
Up-regulated proteins
Down-regulated proteins


Figure S2

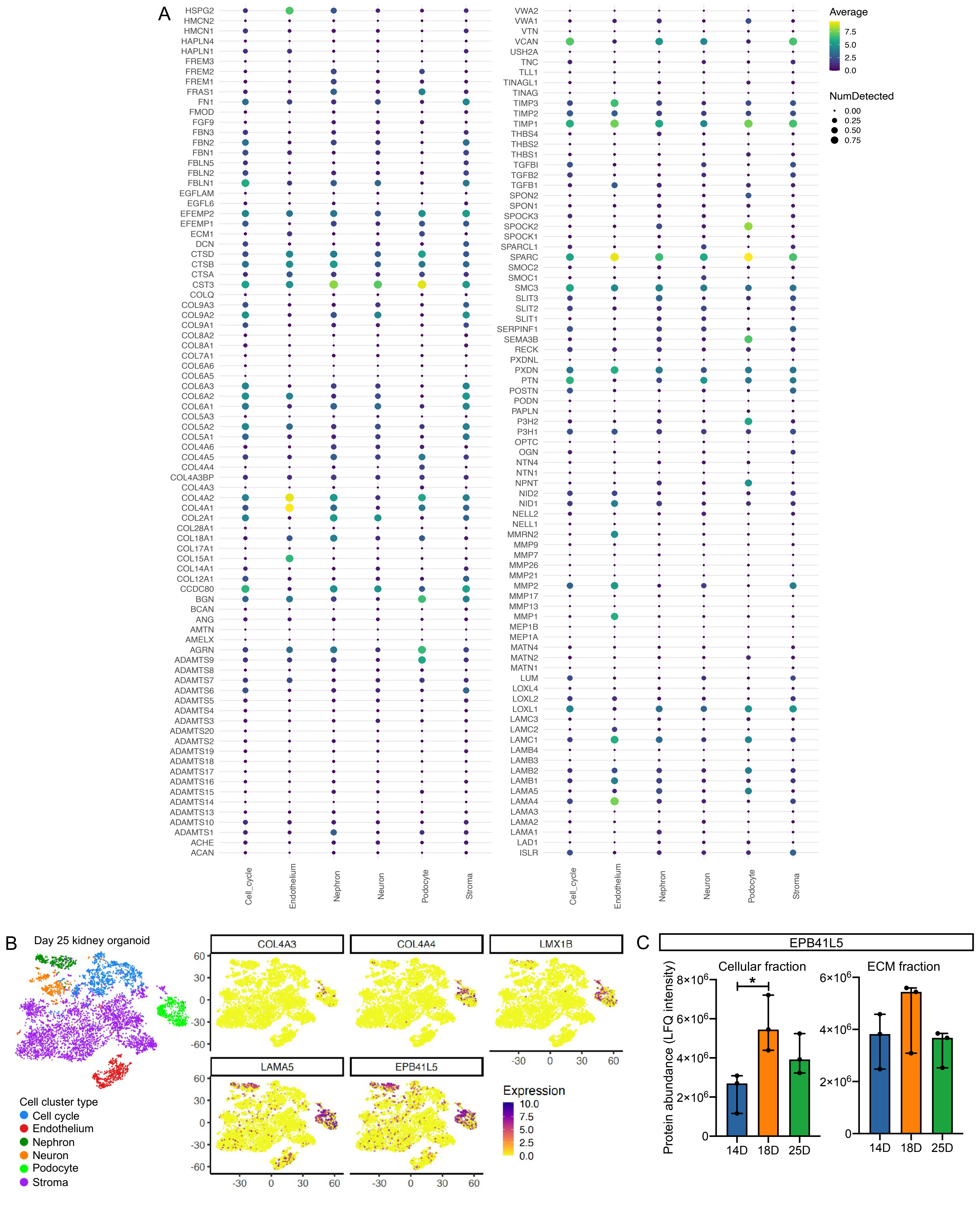

A
HSPG2
HMCN2
HMCN1
HAPLN4
HAPLN1
FREM3
FREM2
FREM1
FRAS1
FN1
FMOD
FGF9
FBN3
FBN2
FBN1
FBLN5
FBLN2
FBLN1
EGFLAM
EGFL6
EFEMP2
EFEMP1
ECM1
DCN
CTSD
CTSB
CTSA
CST3
COLQ
COL9A3
COL9A2
COL9A1
COL8A2
COL8A1
COL7A1
COL6A6
COL6A5
COL6A3
COL6A2
COL6A1
COL5A3
COL5A2
COL5A1
COL4A6
COL4A5
COL4A4
COL4A3BP
COL4A3
COL4A2
COL4A1
COL2A1
COL28A1
COL18A1
COL17A1
COL15A1
COL14A1
COL12A1
CCDC80
BGN
BCAN
ANG
AMTN
AMELX
AGRN
ADAMTS9
ADAMTS8
ADAMTS7
ADAMTS6
ADAMTS5
ADAMTS4
ADAMTS3
ADAMTS20
ADAMTS2
ADAMTS19
ADAMTS18
ADAMTS17
ADAMTS16
ADAMTS15
ADAMTS14
ADAMTS13
ADAMTS10
ADAMTS1
ACHE
ACAN
Cell_cycle
Endothelium
Nephron
Neuron
Podocyte
Stroma
VWA2
VWA1
VTN
VCAN
USH2A
TNC
TLL1
TINAGL1
TINAG
TIMP3
TIMP2
TIMP1
THBS4
THBS2
THBS1
TGFBI
TGFB2
TGFB1
SPON2
SPON1
SPOCK3
SPOCK2
SPOCK1
SPARCL1
SPARC
SMOC2
SMOC1
SMC3
SLIT3
SLIT2
SLIT1
SERPINF1
SEMA3B
RECK
PXDNL
PXDN
PTN
POSTN
PODN
PAPLN
P3H2
P3H1
OPTC
OGN
NTN4
NTN1
NPNT
NID2
NID1
NELL2
NELL1
MMRN2
MMP9
MMP7
MMP26
MMP21
MMP2
MMP17
MMP13
MMP1
MEP1B
MEP1A
MATN4
MATN2
MATN1
LUM
LOXL4
LOXL2
LOXL1
LAMC3
LAMC2
LAMC1
LAMB4
LAMB3
LAMB2
LAMB1
LAMA5
LAMA4
LAMA3
LAMA2
LAMA1
LAD1
ISLR
Cell_cycle
Endothelium
Nephron
Neuron
Podocyte
Stroma
Average
7.5
5.0
2.5
0.0
NumDetected
0.00
0.25
0.50
0.75
B
Day 25 kidney organoid
Cell cluster type
Cell cycle
Endothelium
Nephron
Neuron
Podocyte
Stroma
COL4A3
COL4A4
LMX1B
LAMA5
EPB41L5
60
30
0
-30
-60
-30
0
30
60
Expression
10.0
7.5
5.0
2.5
0.0
C
EPB41L5
Cellular fraction
ECM fraction
Protein abundance (LFQ intensity)
8×10^6
6×10^6
4×10^6
2×10^6
0
*
14D
18D
25D


Figure S3

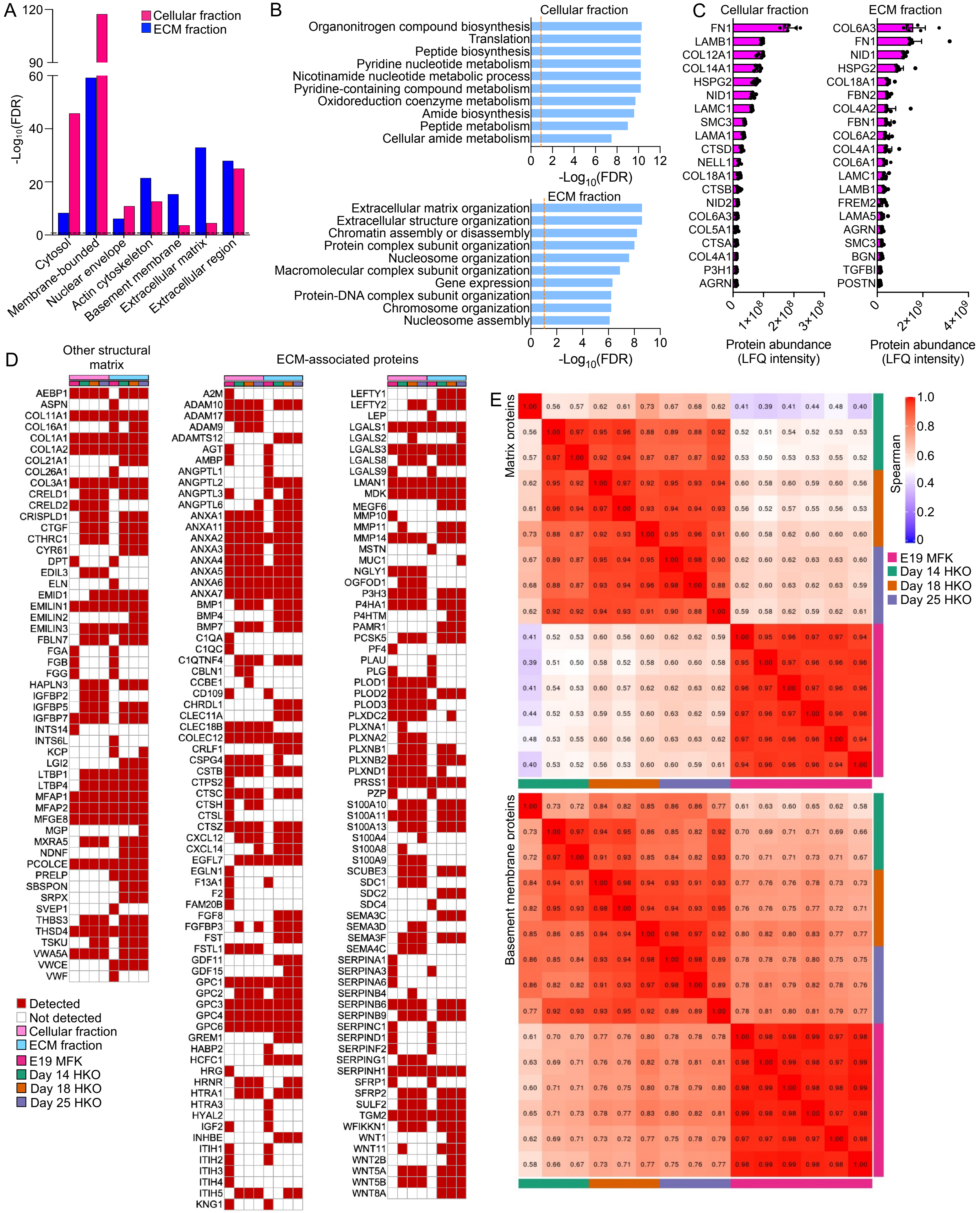
A
Cellular fraction
ECM fraction
-Log10(FDR)
Cytosol
Membrane-bounded
Nuclear envelope
Actin cytoskeleton
Basement membrane
Extracellular matrix
Extracellular region
B
Cellular fraction
Organonitrogen compound biosynthesis
Translation
Peptide biosynthesis
Pyridine nucleotide metabolism
Nicotinamide nucleotide metabolic process
Pyridine-containing compound metabolism
Oxidoreduction coenzyme metabolism
Amide biosynthesis
Peptide metabolism
Cellular amide metabolism
-Log10(FDR)
ECM fraction
Extracellular matrix organization
Extracellular structure organization
Chromatin assembly or disassembly
Protein complex subunit organization
Nucleosome organization
Macromolecular complex subunit organization
Gene expression
Protein-DNA complex subunit organization
Chromosome organization
Nucleosome assembly
-Log10(FDR)
C
Cellular fraction
ECM fraction
Protein abundance (LFQ intensity)
Protein abundance (LFQ intensity)
D
Other structural matrix
ECM-associated proteins
Detected
Not detected
Cellular fraction
ECM fraction
E19 MFK
Day 14 HKO
Day 18 HKO
Day 25 HKO
E
Matrix proteins
Basement membrane proteins
Spearman
E19 MFK
Day 14 HKO
Day 18 HKO
Day 25 HKO

Figure S4

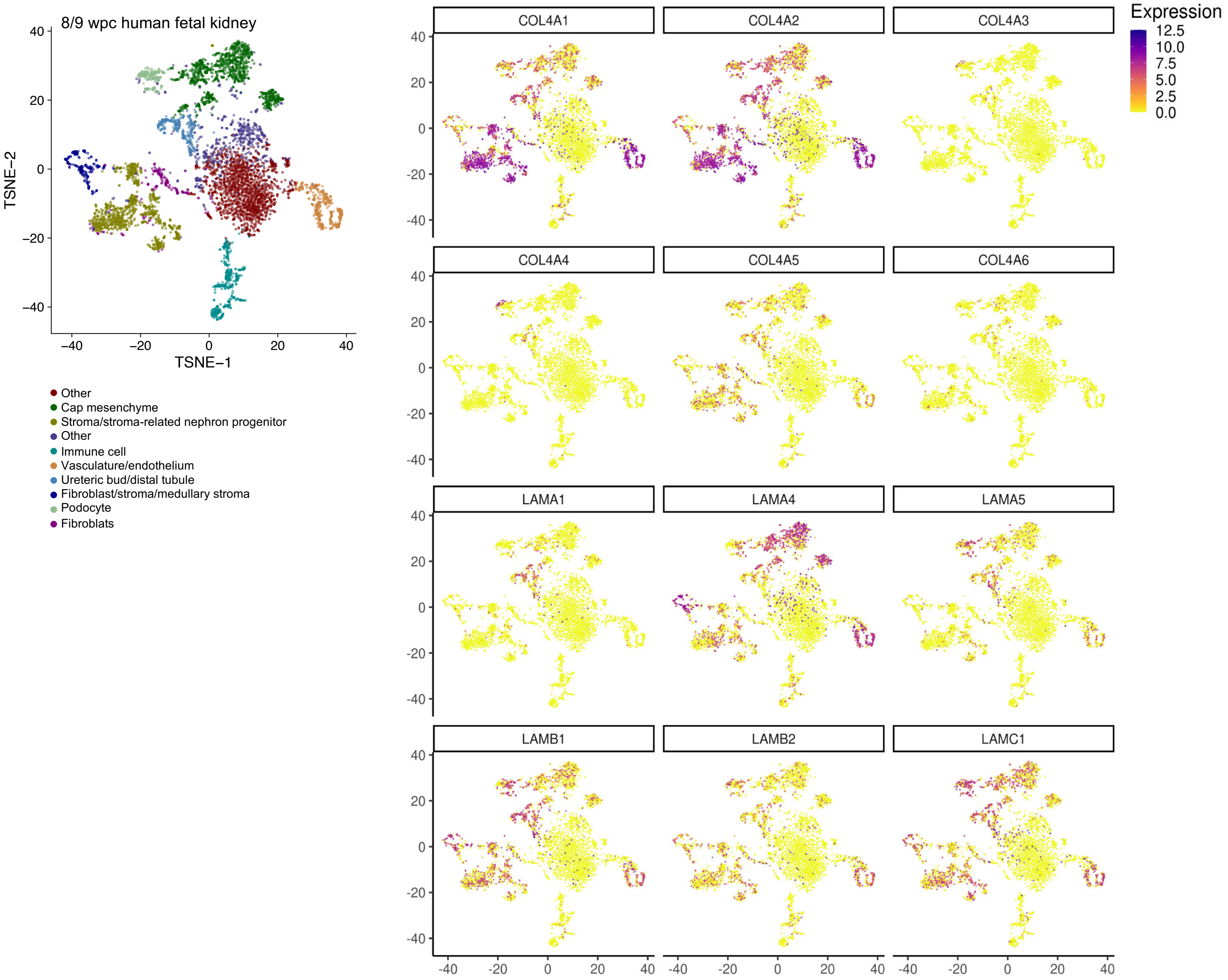
8/9 wpc human fetal kidney
TSNE−1
TSNE−2
Other
Cap mesenchyme
Stroma/stroma-related nephron progenitor
Other
Immune cell
Vasculature/endothelium
Ureteric bud/distal tubule
Fibroblast/stroma/medullary stroma
Podocyte
Fibroblats
COL4A1
COL4A2
COL4A3
COL4A4
COL4A5
COL4A6
LAMA1
LAMA4
LAMA5
LAMB1
LAMB2
LAMC1
Expression
12.5
10.0
7.5
5.0
2.5
0.0

Figure S5

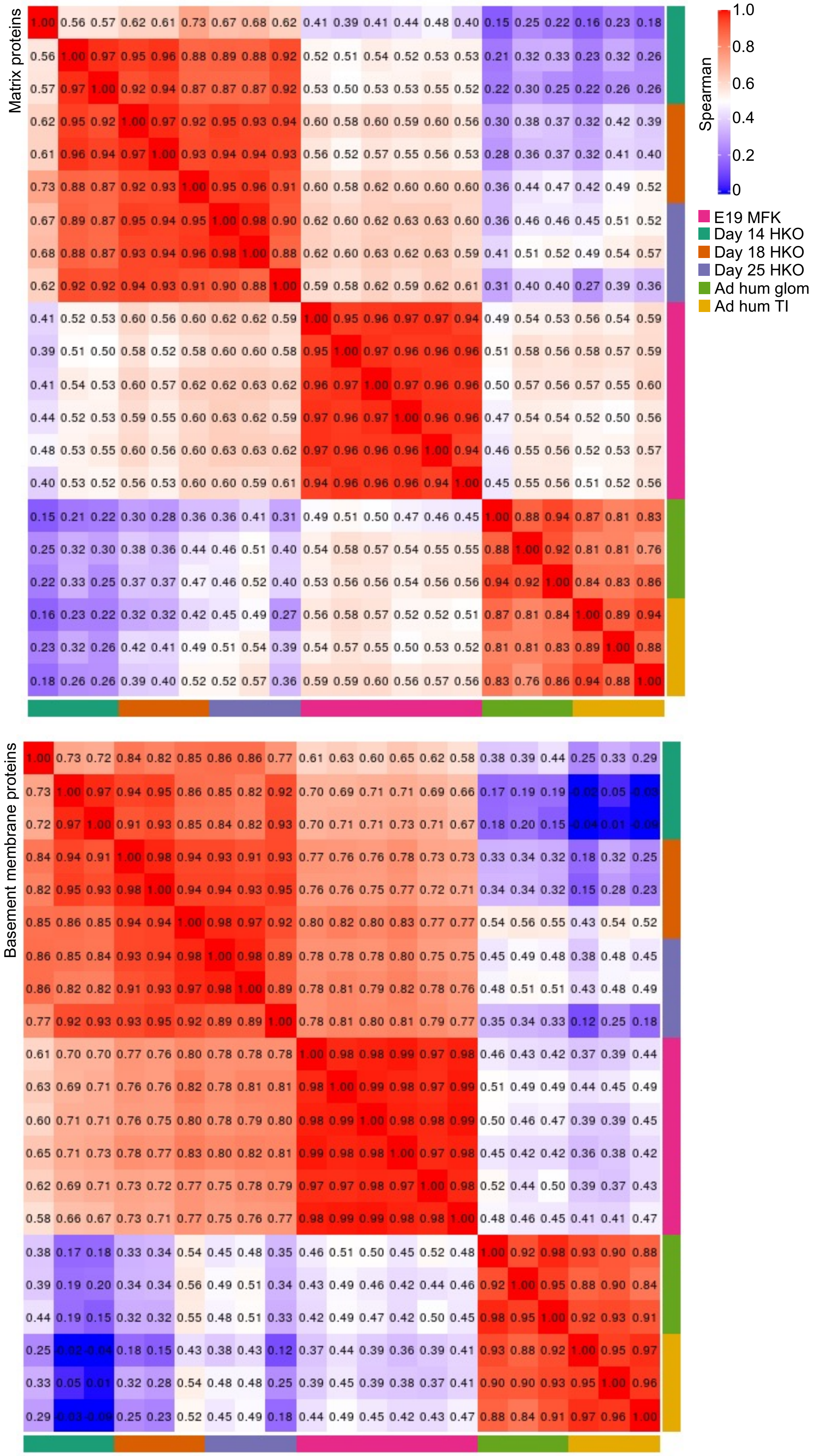
Matrix proteins
Basement membrane proteins
Spearman
1.0
0.8
0.6
0.4
0.2
0
E19 MFK
Day 14 HKO
Day 18 HKO
Day 25 HKO
Ad hum glom
Ad hum TI

Figure S6
